# A Multi-Property Optimizing Generative Adversarial Network for de novo Antimicrobial Peptide Design

**DOI:** 10.1101/2024.11.13.623386

**Authors:** Jiaming Liu, Tao Cui, Tao Wang, Xi Zeng, Yinbo Niu, Shaoqing Jiao, Dazhi Lu, Jun Wang, Shuyuan Xiao, Dongna Xie, Xuecheng Wang, Yongtian Wang, Xuequn Shang, Zhongyu Wei, Jiajie Peng

## Abstract

Antimicrobial peptides (AMPs) play a crucial role in developing novel antiinfective drugs due to their broad-spectrum antimicrobial activity and lower likelihood of causing bacterial resistance. However, laboratory synthesis of AMPs is tedious and time-consuming. Existing computational methods have limited capability in optimizing multiple desired properties simultaneously. Here, we propose a Multi-Property Optimizing Generative Adversarial Network (MPOGAN), a feedback-loop framework that iteratively learns from data with multiple desired properties. This approach enables de novo design of AMPs with potent antimicrobial activity, reduced cytotoxicity, and diversity. Through extensive computational tests, MPOGAN exhibits superior performance in optimizing multiple desired properties of generated AMPs. Ten of the most promising candidates are chemically synthesized, with nine showing potent antimicrobial activity against three bacterial strains and low cytotoxicity against eukaryotic cells. MPOGAN offers a powerful computational approach for effective multi-property optimization of AMPs, thereby advancing the field of AI-aided drug discovery.

## 1 Introduction

As underlined by the World Health Organization’s (WHO) caution, we are rapidly approaching a time when antibiotics may cease to be effective in treating infections [1]. Antimicrobial-resistant pathogens caused more than 2.8 million infections and over 35,000 deaths annually in the United States from 2012 to 2017 [2]. Increasing drug resistance of common pathogens urgently needs the discovery of new effective molecules [3]. Antimicrobial peptides (AMPs), small amphipathic peptides with a net positive charge under physiological pH [4], are often hailed as “natural antibiotics” due to their broad-spectrum antimicrobial activity, minimal toxicity, and lower likelihood of triggering bacterial resistance [5–8]. Therefore, AMPs emerge as potential game-changers in addressing this problem [9].

Discovery of AMPs in the biological laboratory is typically time-consuming and costly, which has driven the study of efficient computational approaches for AMP design [10–12]. One type of widely studied method is predictive models, which take peptides as input and predict specific properties (Antimicrobial activity [13–17], toxicity [18], secondary structure [19], etc.) of these peptides. Employing predictive models trained on natural AMPs, some studies develop the quantitative structure-activity relationship (QSAR) models to identify potential AMPs in a peptide database that have not been experimentally validated [4, 20, 21]. These predictive models can identify existing peptides with desired properties, but cannot design novel AMPs. Another type of computational method is genetic algorithms [22, 23], which guide peptide sequence evolution by randomly mutating existing AMPs and viability assessment. However, these methods are limited by the local search space, making it difficult to discover potential AMPs that are significantly different from known AMPs but are more effective. Recently, deep generative methods in machine learning have emerged as efficient computational approaches to generate AMPs de novo. These methods frequently employ Autoregressive Models (AMs) [24–27], Variational Autoencoders (VAEs) [10, 11, 28–30], and Generative Adversarial Networks (GANs) [7, 31–36]. However, these methods cannot optimize multiple properties of AMPs.

The clinical application of AMPs requires potent antimicrobial activity and limited side effects. One prominent side effect is the cytotoxicity against eukaryotic cells, which is a major obstacle to AMPs becoming therapeutic agents [6, 18, 36, 37]. Unfortunately, existing methods focus primarily on optimizing antimicrobial activity, struggling with generating AMPs with multiple desired properties. Although some approaches [4, 10, 11, 38] have implemented additional in silico step-by-step screening to identify AMP candidates with multiple desired properties, the success rate of AMP wet-laboratory verification remains constrained. Therefore, developing a method for multi-property optimization in de novo AMP design is imperative. Essentially, the antimicrobial activity and cytotoxicity of AMPs are influenced by their key physicochemical features, such as net charge and amphiphilicity [10, 39]. However, it is challenging to optimize physicochemical features simultaneously, due to their inherent conflict. For example, increasing net charge or hydrophobicity could boost antimicrobial activity, but it concurrently increases the risk of toxicity to normal cells [39]. In this context, a straightforward idea is to train a model using AMPs with multiple desired properties. Unfortunately, for most experimentally validated AMPs, only the antimicrobial activity validation is performed, resulting in the lack of training data for multi-property optimization [40].

This study introduces a Multi-Property Optimizing Generative Adversarial Network (MPOGAN). We propose an extended feedback-loop GAN framework, specifically designed to tackle the scarcity of high-quality AMPs with multiple desired properties as training data. To fill this gap, multiple plug-and-play model-embedded evaluators are designed to screen for high-quality training data from generated peptides. Furthermore, a Real-Time Knowledge-Updating (RTKU) strategy is proposed to iteratively update the training dataset dynamically. By learning from iteratively generated training datasets, MPOGAN can design novel AMPs with potent antimicrobial activity, reduced cytotoxicity, and diversity. The extensive evaluation of AMP candidates indicates that MPOGAN outperforms other state-of-the-art methods [10, 24, 26, 28, 30]. Ten of the most promising predicted candidates are chemically synthesized. Among these, nine exhibited potent antimicrobial activity against diverse Gram-positive and Gram-negative pathogens, while demonstrating low cytotoxicity against MC3T3-E1 cells. In addition, MPOGAN achieved a high success rate of 90% in AMP wet-laboratory validation, surpassing baseline methods under severe restrictions. In summary, MPOGAN offers a powerful computational approach for de novo design of AMPs with multiple desired properties, thereby advancing AI-aided drug discovery.

## 2 Results

### 2.1 MPOGAN: a deep generative framework for de novo AMP design

To de novo design AMPs with multiple desired properties, including potent antimicrobial activity, reduced cytotoxicity, and diversity, we propose the MPOGAN framework (Fig. 1), which consists of two main stages: a pre-training stage (Fig. 1a left) and a multi-property optimizing (MPO) stage (Fig. 1a right). In the pre-training stage, MPOGAN undergoes generator pre-training, discriminator pre-training, and adversarial learning using experimentally validated AMPs and non-AMPs. This process enables MPOGAN to learn the characteristics of real AMPs and generate analogous peptides. Inspired by SeqGAN [41], MPOGAN consists of a generator (Fig. 1b) based on a recurrent neural network (RNN) and a discriminator (Fig. 1c) based on a protein large language model (ProLLM). The generator produces amino acid sequences from an initial input token. By modeling the generator with an RNN, MPOGAN can learn the semantic relationships between discrete amino acids. The discriminator consists of a Protein LLM called ESM-2 [42] for feature encoding, and a Multilayer Perceptron (MLP) for binary classification based on these representations. The discriminator takes amino acid sequences as input and outputs the probabilities of being AMPs. Using ProLLM for feature encoding enables MPOGAN to leverage the inherent biological information between amino acids [36].

**Fig. 1.**
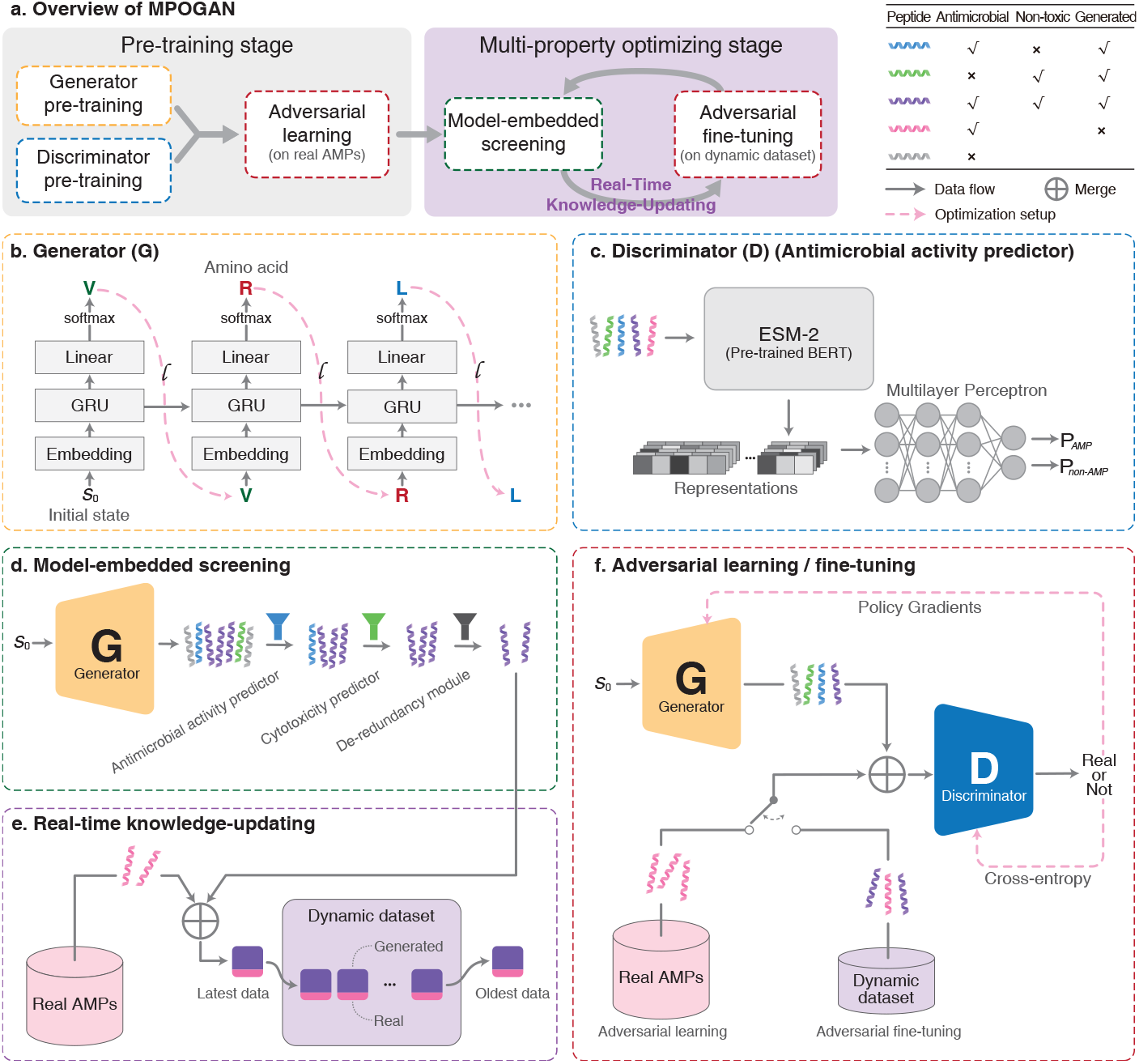
The workflow of MPOGAN. **a** Overview of MPOGAN. The pre-training stage (left) contains generator pre-training (yellow), discriminator pre-training (blue), and adversarial learning on real AMPs (red); the multi-property optimizing (MPO) stage (right) consists of multiple iterations of model-embedded screening (green), real-time knowledge-updating (purple), and adversarial finetuning on dynamic dataset (red). The color of the dashed box corresponds to b-f. **b** The generator is based on a Recurrent Neural Network (RNN) consisting of three layers. **c** The discriminator consists of a pre-trained Bidirectional Encoder Representations from Transformers (BERT) (ESM-2 [42]) with fixed parameters and a Multilayer Perceptron (MLP) with trainable parameters. The antimicrobial activity predictor shares the same architecture with the discriminator. **d** A model-embedded screening process comprises three plug-and-play evaluators (funnels), where the parameters of all evaluators are fixed. **e** A real-time knowledge-updating strategy updates the dynamic dataset via first-in-first-out, where the dynamic dataset has a fixed size. **f** Adversarial learning on real AMPs during the pre-training stage, and adversarial fine-tuning on the dynamic dataset during the MPO stage.

In the MPO stage, we enhance MPOGAN with the ability to optimize multiple properties through a model-embedded screening process (Fig. 1d), a Real-Time Knowledge-Updating (RTKU) strategy (Fig. 1e), and adversarial fine-tuning of MPOGAN (Fig. 1f). Firstly, to screen high-quality candidates from MPOGAN-generated peptides, the model-embedded screening includes three plug-and-play evaluators: a ProLLM-based antimicrobial activity predictor (LLM-AAP) trained on the real AMP dataset to identify highly potent AMP candidates, a cytotoxicity predictor [18] to select candidates with reduced cytotoxicity, and a de-redundancy module [43] to ensure diversity among the candidates. Then, following the feedback-loop architecture [33], we propose the RTKU strategy to guide the updating of a dynamic dataset. The data used for updating consists of two parts: a large portion of high-quality candidates from the model-embedded screening, which guides MPOGAN to learn high-quality features, and a small portion from the real AMP dataset, which ensures the generalization capability of MPOGAN. The RTKU strategy ensures MPOGAN captures new features without ignoring the existing ones. Finally, adversarial fine-tuning is performed to optimize MPOGAN using the dynamic dataset, thereby fully leveraging the knowledge acquired by model-embedded screening. Further details about the MPOGAN framework can be found in the “Methods” section.

### 2.2 MPOGAN outperforms the baseline methods in generating AMPs de novo

We compared the performance of the MPOGAN with five state-of-the-art deep generative methods in generating high-quality AMPs: Dean-VAE [30], Nagajaran-LSTM [24], PepCVAE [28], HydrAMP [10], and Muller-RNN [26]. Following the evaluation protocols provided by Szymczak et. al. [10], we mainly evaluated three properties of generated AMPs, including antimicrobial activity, cytotoxicity, and diversity. The details of comparison models and evaluation settings can be found in Supplementary Note 2.1.

Compared to other methods, the median probability of antimicrobial activity for MPOGAN is 99.99%, which achieves the best performance (Fig. 2a); and the median probability of cytotoxicity is 64.70%, which is 2% lower than the existing state-of-the-art method (Muller-RNN) (Fig. 2b). For diversity evaluation, MPOGAN has 99.44% of unique generated sequences, which is 1.03% higher than the second highest (Nagajaran-LSTM) (Fig. 2c). The proportion of peptides considered to have potent antimicrobial activity is 88.32% (Fig. 2d), which is 3.72% higher than the second highest; the proportion of peptides with reduced cytotoxicity is 75.60%, which is 11.25% higher than the second lowest (Fig. 2e); and the proportion of high-quality peptides (simultaneously meeting the requirements of potent antimicrobial activity and reduced cytotoxicity) is 66.43%, which is 15.86% higher than the second highest (Fig. 2f). The results demonstrate that MPOGAN outperforms baseline methods in generating AMP candidates with each of the three properties. Furthermore, MPOGAN significantly surpassed existing baseline methods in generating AMPs that efficiently balance potent antimicrobial activity with reduced cytotoxicity. While some baseline methods (Nagajaran-LSTM, PepCVAE, and HydrAMP) can generate AMPs with potent antimicrobial activity, the cytotoxicity profile of AMP candidates is not ideal due to the lack of targeted optimization. In conclusion, MPOGAN outperforms existing state-of-the-art methods in generating high-quality AMP candidates de novo.

**Fig. 2.**
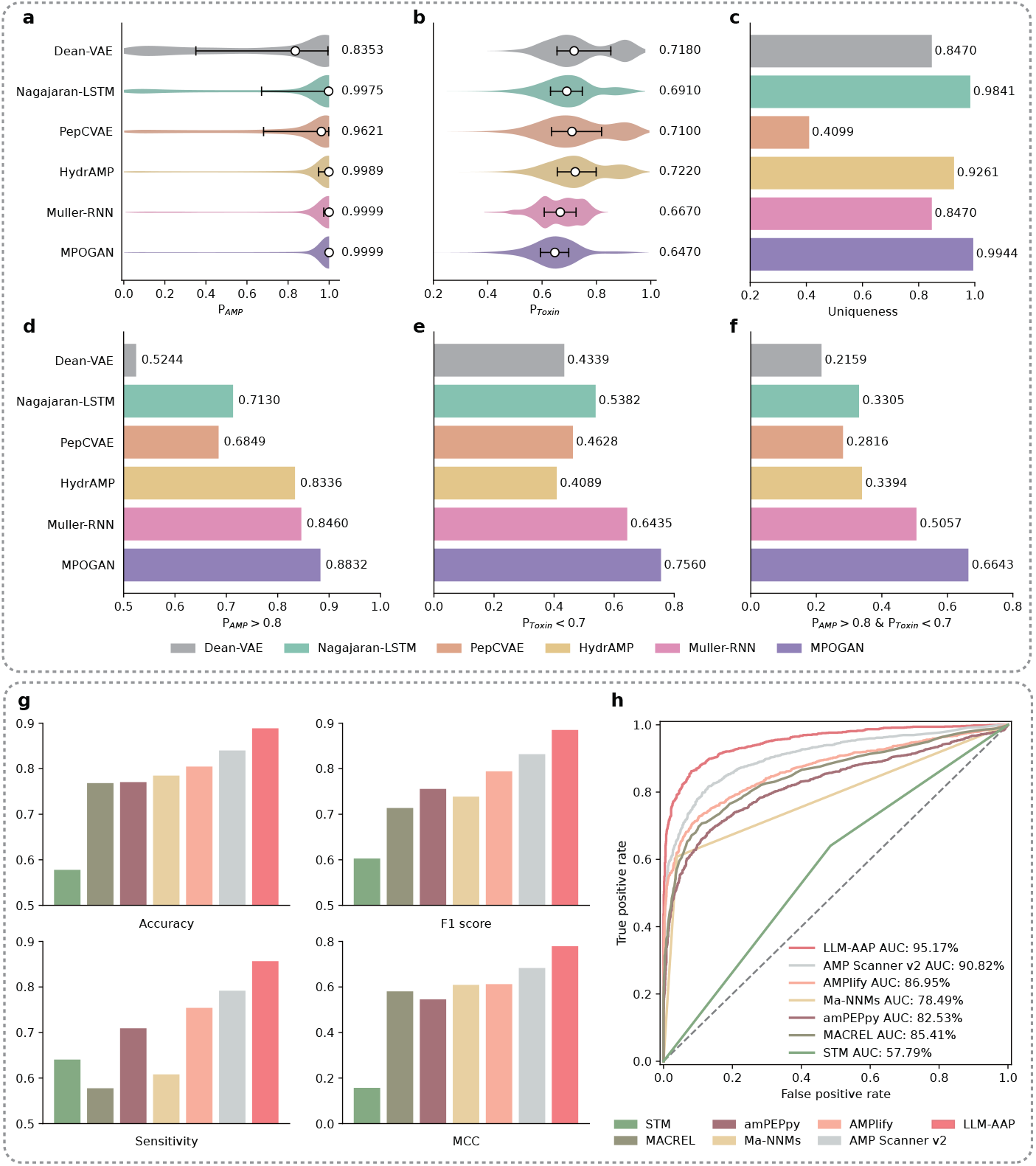
Comparison assessments of MPOGAN and generative baseline methods on generating AMPs de novo, as well as LLM-AAP and predictive baseline methods on antimicrobial activity prediction. **a-f** Generative performance of MPOGAN, in comparison to Dean-VAE, Nagajaran-LSTM, PepCVAE, HydrAMP, and Muller-RNN. Evaluations were performed on generated AMP candidates. Antimicrobial activity was evaluated by trained LLM-AAP, and cytotoxicity was evaluated by Toxinpred2 [18]. **a, b** The probability distribution of **a** antimicrobial activity score (*P*_AMP_) and **b** cytotoxicity score (*P*_Toxin_). The white dots and values on the right mark the median of each distribution, and the black horizontal lines denote the interquartile range of each distribution. **c** The proportion of unique peptides. **d-f** The proportion of peptides with **d** potent antimicrobial activity peptides, **e** low cytotoxicity, and **f** both of them. **g, h** Predictive performance oef LLM-AAP, in comparison to STM, MACREL, amPEPpy, Ma-NNMs, AMPlify, and AMPScanner v2 on the test set. **g** Accuracy, F1 score, sensitivity, and Matthews correlation coefficient (MCC) of predictive models. **h** Receiver-operating characteristic (ROC) curves of predictive models. The diagonal dashed line indicates random prediction.

### 2.3 Performance evaluation of antimicrobial activity predictor

Antimicrobial activity prediction can be regarded as an AMP identification task. We compared the performance of our ProLLM-based Antimicrobial Activity Predictor (LLM-AAP) with six state-of-the-art AMP identification methods: amPEPpy [14, 44], MACREL [13], Ma-NNMs [20], AMPlify [17], AMP Scanner v2 [16], and STM [15].

We applied accuracy, sensitivity, F1 score, Matthews Correlation Coefficient (MCC), and Area Under the Curve (AUC) as the model evaluation metrics. Details of these methods and evaluation settings can be found in Supplementary Note 2.2.

The experiment results show that LLM-AAP achieves the highest performance among all methods in five evaluation metrics (Fig. 2g, h, Supplementary Table 1, and Supplementary Fig. 4). Compared to other methods, the accuracy of our predictor is 4.87% higher than the runner-up methods (from 0.8400 to 0.8887), the F1 score is 5.31% higher (from 0.8319 to 0.8850), the sensitivity is 6.49% higher (from 0.7917 to 0.8566), the MCC is 9.57% higher (from 0.6832 to 0.7790), and the AUC is 4.35% higher (from 90.82% to 95.17%). The improvement in results is attributed to the fact that LLM-AAP enables the extraction of complex intrinsic biological information from peptides using ProLLM. In conclusion, LLM-AAP can achieve significant improvements in identifying AMP candidates compared to state-of-the-art methods.

The RTKU strategy of MPOGAN updates the AMP candidates refined by the model-embedded screening into the dynamic dataset. Therefore, the quality of these peptides directly affects the optimization direction for MPOGAN. If the model-embedded screening cannot accurately identify the desired AMP candidates, MPOGAN will be fine-tuned in an unknown direction. As the first evaluator of the model-embedded screening process, the superior performance of LLM-AAP ensures that MPOGAN continuously learns the features of AMPs with potent antimicrobial activity during adversarial fine-tuning. Additionally, the MPOGAN discriminator has the same network architecture as LLM-AAP, ensuring that the superior performance of LLM-AAP translates to a highly effective MPOGAN discriminator with robust discrimination ability.

### 2.4 Features of AMP candidates generated by MPOGAN

In this section, we evaluated the distributions of key molecular features implicated in the antimicrobial nature of MPOGAN-generated AMPs (MPOPs). Here, we compared the key molecular features of three sets, including the experimentally validated AMPs we collected (labeled as real AMPs), the 50,000 AMP candidates directly generated by MPOGAN (labeled as MPOPs-50k), and the 124 stringent AMP candidates (labeled as MPOPs-124). MPOPs-124 includes the most promising AMP candidates from MPOPs-50k. Details on generating MPOPs-124 can be found in “Methods” and Supplementary Fig. 7. The key molecular features analyzed include amino acid composition, charge, isoelectric point (pI), aromaticity, Eisenberg hydrophobicity, hydrophobic moment, hydrophobic ratio, charge density, instability index, and aliphatic index. These molecular features are implicated in the peptide-membrane interaction and structural stability [11, 45].

Compared to real AMPs, both MPOPs-50k and MPOPs-124 contain higher proportions of Phenylalanine (F) and Isoleucine (I), which have hydrophobic side chains; higher proportions of Histidine (H) and Lysine (K), which have positively charged side chains; and lower proportions of Aspartic acid (D) and Glutamic acid (E), which have negatively charged side chains (Fig. 3a). These results demonstrate that MPOGAN can generate AMP candidates with enhanced lipophilicity and a stronger positive net charge. These two features help peptides better penetrate and disrupt the cell membrane of bacteria [8]. The MPOPs-124 contains a higher proportion of neutral amino acids (Methionine (M), Asparagine (N), Glutamine (Q), Threonine (T), Serine (S)) compared to both the real AMPs dataset and MPOPs-50k, as well as fewer Tryptophan (W), Arginine (R), and Proline (P) (Fig. 3a). This indicates that MPOPs-124 peptides exhibit a more balanced hydrophilic profile, potentially enhancing their solubility and stability in aqueous environments. Additionally, the reduction in W, R, and P might contribute to a lower likelihood of aggregation and structural rigidity, thereby improving their flexibility and ability to adapt to different bacterial membrane environments. The overall amino acid composition of MPOPs-124 suggests a subtle optimization for antimicrobial activity, combining hydrophilic balance with enhanced peptide-membrane interaction.

**Fig. 3.**
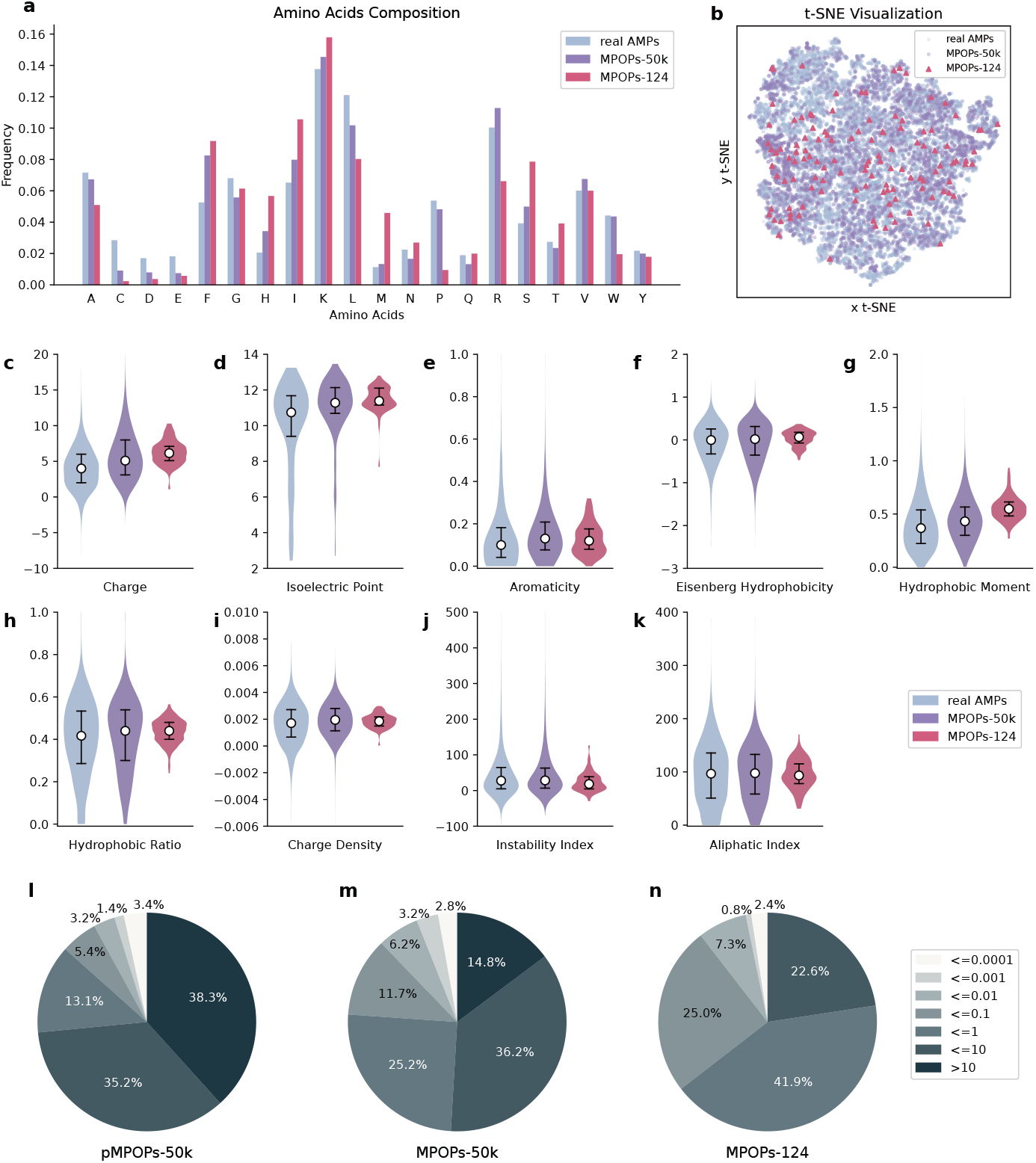
Comparison of the physicochemical nature of real AMPs and MPOGAN-generated AMP candidates. **a, c-k** Comparison of **a** amino acid composition, **c** charge, **d** isoelectric point (pI), **e** aromaticity, **f** Eisenberg hydrophobicity, **g** hydrophobic moment, **h** hydrophobic ratio, **i** charge density, **j** instability index, and **k** aliphatic index distribution of real AMPs with MPOGAN-generated AMP candidates (MPOPs-50k: 50,000 peptides that are directly generated by MPOGAN; MPOPs-124: 124 peptides that are obtained through preliminary selection process). All physicochemical properties are calculated using modlAMP [70]. **b** t-SNE visualization. 9 physicochemical features of peptides are calculated and downscaled to 2 dimensions with a perplexity of 30. **l-n** The Expect value (E-value) distribution of pMPOPs-50k (50,000 peptides that are directly generated by **l** pre-trained MPOGAN), **m** MPOPs-50k, and **n** MPOPs-124.

As shown in Fig. 3c-k, MPOPs-50k maintain physicochemical features similar to real AMPs, while MPOPs-124 exhibit a more concentrated distribution across various physicochemical features. In detail, for some features, including charge, isoelectric point, and hydrophobic moment, the mean values of MPOPs-50k are higher than those of real AMPs, and the mean values of MPOPs-124 are even higher. The enhancement of these properties is beneficial for peptide-membrane interactions [45]. Additionally, peptides with an instability index (II) below 40 are considered stable [46]. The mean value of II for MPOPs-50k and MPOPs-124 is lower than that of real AMPs. The proportion of peptides with II ¡ 40 in MPOPs-50k (60.57%) increased by 0.5%, and in MPOPs-124 (76.61%) increased by 16.54%, compared to real AMPs (60.07%). This finding indicates that the generated AMP candidates, particularly those in the MPOPs-124, are more stable than real AMPs. We further evaluated the overall distribution of the three datasets across the nine physicochemical properties (Fig. 3b). By reducing the dimensionality of the standardized nine physicochemical properties to two dimensions using t-SNE [47], we observed that the distribution of MPOPs-50k is similar to that of real AMPs, while the distribution of MPOPs-124 is more concentrated, aligning with previous findings. These observations demonstrate that MPOGAN is effective in generating AMP candidates with physicochemical features similar to those found in natural AMPs. The MPOPs-124 refined the candidate pool to focus on peptides with more specific and desired features.

We also evaluated the homology of AMP candidates against real AMPs. As shown in Fig. 3l-n and Supplementary Table 2, we calculated the percentage of pMPOPs-50k (50,000 AMP candidates generated by pre-trained MPOGAN), MPOPs-50k, and MPOPs-124 in different categories of Expect value (E-value) [48]. The E-value for the match with the highest score was considered, as obtained by performing a BLAST similarity search [49] against real AMPs. A larger E-value implies fewer repeated sequence fragments and a smaller likelihood of homology. Compared with pMPOPs-50k, MPOPs-50k exhibits higher homology against real AMPs, and MPOPs-124 exhibits the highest. This finding suggests that the MPO stage improves the correlation between AMP candidates and real AMPs. Meanwhile, the AMP candidate sequences generated by MPOGAN still maintain diversity (Fig. 2c), and the majority of E-values are still higher than 0.01 (87.9% for MPOPs-50k and 89.5% for MPOPs-124), indicating that the MPO stage does not sacrifice sequence diversity while optimizing the molecular properties of AMP candidates. The analysis of sequence alignment (Supplementary Fig. 6) also indicates that AMP candidates generated by MPOGAN display outstanding diversity and novelty.

### 2.5 Quantitative analysis of MPO stage

The main advantage of MPOGAN is that it optimizes multiple desired properties of AMP candidates in the MPO stage. In this stage, the model-embedded screening is employed to obtain high-quality AMP candidates from MPOGAN-generated peptides at each iteration. To comprehensively assess the effectiveness of the MPO stage, we evaluated changes in four AMP candidate sets iteratively obtained in the model-embedded screening throughout the MPO stage: (1) candidates generated by the MPOGAN generator 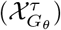, (2) candidates screened by the antimicrobial activity predictor 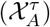, (3) candidates screened by the cytotoxicity predictor 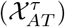, and (4) candidates screened by the de-redundancy module 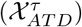. The iteration of MPOGAN optimization was performed for 700 epochs.

Fig. 4a shows the change in the number of AMP candidates passing every evaluator. Specifically, the number of 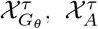, and 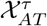 show a gradual downward trend during the iteration, with decreasing descent magnitude, indicating that the MPOGAN is gradually reaching convergence. The number of 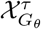 shows the most significant decrease (70.87%), indicating that the time consumption of the MPO stage is effectively controlled. In other words, as the iteration progresses, the MPOGAN generator needs to generate fewer AMP candidates to meet the requirements of the model-embedded screening. Meanwhile, the number of 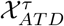 remains stable across all iterations. These observations suggest that the RTKU strategy not only ensures that the amount of candidates meets the requirements for updating the dynamic dataset, but also effectively controls the quantity of 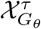, thus maximizing the efficiency of the plug-and-play model-embedded screening.

**Fig. 4.**
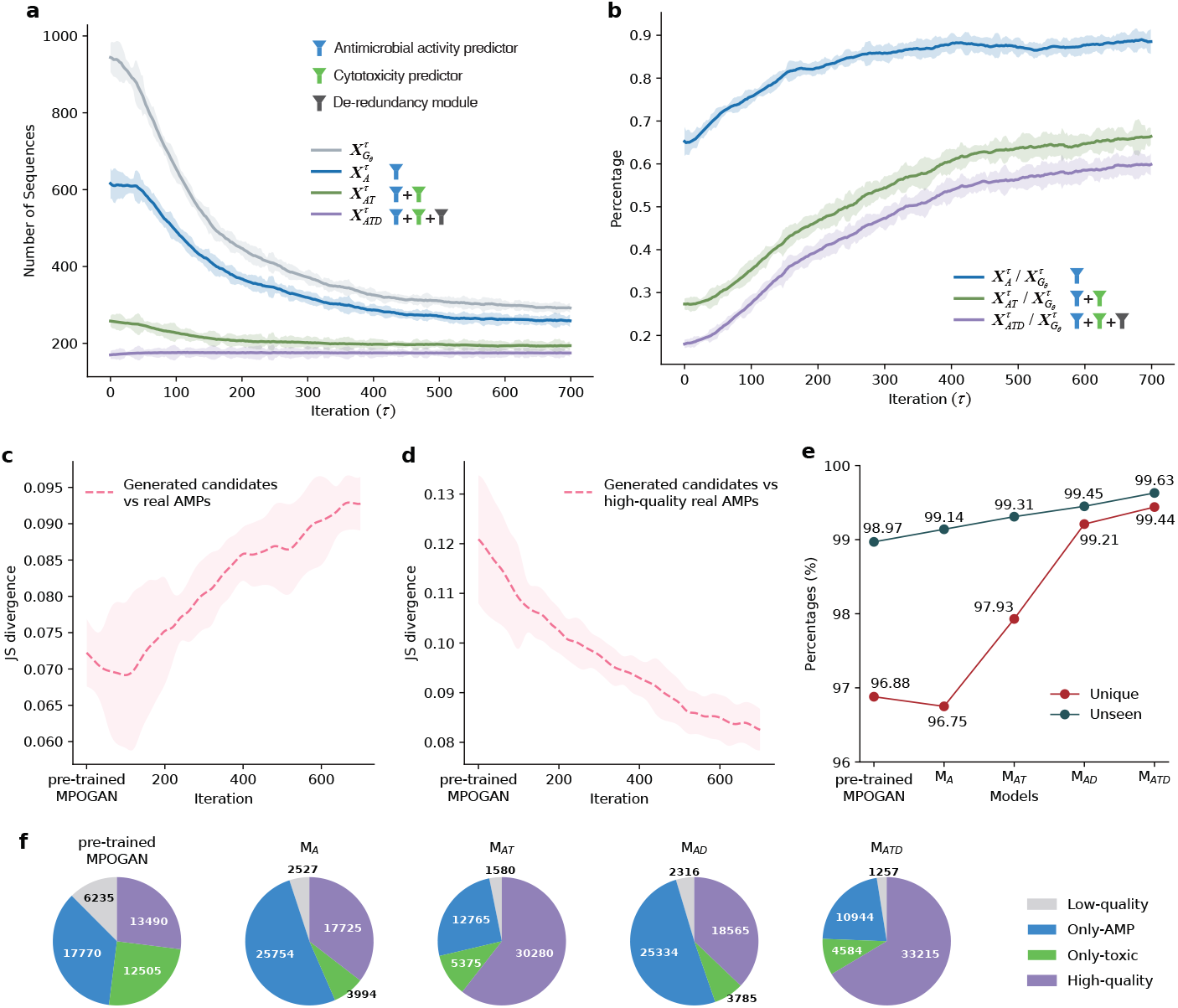
Analysis of MPO stage and ablation studies. **a-d** Analysis of MPO stage. The translucent area outside the curve indicates the range of fluctuations in the data. **a** Variation with iteration (*τ*) in the size of the AMP candidate set generated by the MPOGAN-generator and the size of AMP candidate sets that pass the model-embedded evaluators step-by-step in the model-embedded screening. **b** Percentage variation with *τ* in the size of AMP candidate sets that pass the model-embedded evaluators relative to the size of AMP candidate set generated by the MPOGAN-generator. **c, d** Variation with *τ* in Jensen-Shannon (JS) divergence of the distribution of the AMP candidate set generated by the MPOGAN-generator relative to the distribution of **c** the real AMP dataset, and **d** the high-quality subset in the real AMPs dataset. **e, f** Ablation studies for various combinations of model-embedded evaluators. Each generated set contains 50,000 peptides. **e** Percentage variation in the size of unique and unseen AMP candidate sets relative to the size of the AMP candidate set generated by the generator. **f** Property distribution of AMP candidate sets generated by the generator. The numbers represent the subset size.

To assess the fine-tuning process of MPOGAN during the MPO stage, we analyzed the percentage change of AMP candidates passing each evaluator relative to 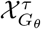 (Fig. 4b). There is a consistent increase in the percentage of sequences passing through different model-embedded evaluators with each iteration. Specifically, the percentage of AMP candidates passing the antimicrobial activity predictor 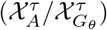 improves by 25.14% (from 62.13% to 87.27%), the cytotoxicity predictor 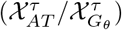 improves by 41.89% (from 27.20% to 69.09%), and the de-redundancy module 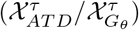 improves by 44.74% (from 18.53% to 63.27%). These observations indicate that the MPO stage has significantly enhanced the performance of MPOGAN in generating multi-property superior AMP candidates. We also observed that, in every iteration, the percentage of 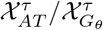 (candidates meeting both antimicrobial activity and cytotoxicity requirements) significantly decreases compared to those before the cytotoxicity screening (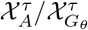, i.e., candidates meeting only the activity requirements), reducing by approximately 35% (before MPO stage) and 18% (after MPO stage). This implies that a substantial portion (approximately 56% (before MPO stage) and 21% (after MPO stage)) of peptides meeting the antimicrobial activity criteria cannot satisfy the cytotoxicity requirements. This phenomenon confirms the necessity of optimizing the cytotoxicity of AMPs in generative methods, and it also demonstrates that the model-embedded screening can effectively remove candidates that do not meet multi-property optimization objectives.

To further assess the robustness of the RTKU strategy, we also analyzed how the data composition of the dynamic training set changes throughout the iterations (Supplementary Note 3 and Supplementary Fig. 3). The results indicate that the generated peptides in the dynamic dataset quickly reach the expected quantity ratio with the real AMPs and then maintain the dynamic equilibrium. In conclusion, the RTKU strategy not only enhances the iterative efficiency but also ensures the stability of knowledge updates.

To strengthen our findings, we further evaluated the changes in the data distribution of AMP candidates generated by MPOGAN during the MPO stage, as compared to the real AMP dataset (Fig. 4c-d and Supplementary Fig. 5). We employed trained models in different iterations to generate AMP candidates, ranging from the 0th iteration (i.e., the pre-trained MPOGAN) to the 700th iteration (i.e., the final MPOGAN). The model parameters are saved every 20 iterations, resulting in a total of 36 models. Each model generates 5000 AMP candidates. We employed ESM-2 (esm2 t6 8M UR50D) [42] to extract feature representations of the real AMPs and all AMP candidates. We then downscaled the feature representations to two dimensions using Principal Component Analysis (PCA) [50]. As shown in Fig. 4c-d, we utilized Jensen-Shannon (JS) divergence [51] to evaluate the similarity between the feature distribution of candidates generated by 36 models in the MPO stage and the real AMPs, as well as the similarity to high-quality real AMPs (i.e., peptides in the real AMPs dataset that satisfy *P*_AMP_ *>* 0.8 & *P*_Toxin_ *<* 0.7). The smaller the value of JS divergence (ranging from [0, 1]), the closer the two distributions are. As the iterations progressed, the similarity between MPOGAN-generated candidates and the real AMPs gradually decreased, while the similarity to high-quality true AMPs gradually increased. These observations indicate that MPOGAN can effectively utilize high-quality peptide information during fine-tuning in the MPO stage, and disregard information that resembles real AMPs but does not meet multi-property requirements.

### 2.6 Ablation studies on MPOGAN

The optimizing direction and performance of MPOGAN largely depend on the model-embedded evaluators. To validate the effectiveness of the three model-embedded evaluators (antimicrobial activity predictor (*f*_AMP_), cytotoxicity predictor (*f*_Toxin_), and de-redundancy module (*f*_DR_)), we conducted comprehensive ablation studies. Specifically, we evaluated the impact of five evaluator combinations on the performance of MPOGAN. These combinations and their corresponding generative models include: (1) Pre-trained MPOGAN without any evaluators; (2) Only *f*_AMP_ (*M*_*A*_); (3) Combines *f*_AMP_ and *f*_Toxin_ (*M*_*AT*_); (4) Combines *f*_AMP_ and *f*_DR_ (*M*_*AD*_); (5) Combines *f*_AMP_, *f*_Toxin_ and *f*_DR_ (*M*_*ATD*_, i.e., the final MPOGAN). All models were trained uniformly using identical hyperparameters, and each generated 50,000 AMP candidates for evaluation. Fig. 4e shows the percentages of AMP candidates that are non-repetitive (*Unique*) and those not appearing in the real AMP dataset (*Unseen*) for the five combinations. *M*_*ATD*_ achieved the best *Unique* and *Unseen* value. This indicates that the diversity and novelty of the AMP candidates are significantly improved during the MPO stage. We also observe that although *M*_*A*_ has a *Unique* value of 0.13% lower than pre-trained MPOGAN, its *Unseen* value increased by 0.16%. This suggests that optimizing antimicrobial activity may slightly reduce the diversity of AMP candidates, while increasing their novelty. Furthermore, the *Unique* values of *M*_*AT*_, *M*_*AD*_, and *M*_*ATD*_ have significantly increased compared to pre-trained MPOGAN and *M*_*A*_. This indicates that the optimization of cytotoxicity and/or diversity completely offsets the negative impact of antimicrobial activity optimization on the diversity of AMP candidates. This once again highlights the importance of cytotoxicity optimization in AMP design.

We further compared the compositions of AMP candidates generated by the models corresponding to the five evaluator combinations mentioned above. We utilized LLM-AAP and Toxinpred2 [18] to score these candidates, dividing them into four non-overlapping groups: (1) Candidates that meet *P*_AMP_ ≤ 0.8 & *P*_Toxin_ ≥ 0.7, considered to neither satisfy antimicrobial activity nor cytotoxicity requirements (*Low-quality* group); (2) Candidates that meet *P*_AMP_ *>* 0.8 & *P*_Toxin_ ≥ 0.7, considered only to satisfy antimicrobial activity requirements, but not cytotoxicity (*Only-AMP* group); (3) Candidates that meet *P*_AMP_ ≤ 0.8 & *P*_Toxin_ *<* 0.7, considered only to satisfy cytotoxicity requirements, but not antimicrobial activity (*Only-toxin* group); (4) Candidates that meet *P*_AMP_ *>* 0.8 & *P*_Toxin_ *<* 0.7, considered to satisfy both antimicrobial activity and cytotoxicity requirements simultaneously (*High-quality* group). Our goal is to obtain a larger proportion of the *High-quality* group. Fig. 4f presents the composition of these four groups of AMP candidates generated by the five evaluator combinations, where *M*_*ATD*_ achieves the best performance in the proportion of the *High-quality* group. Compared to pre-trained MPOGAN, the *High-quality* group in *M*_*A*_ increased by 31.39%, and the *Only-AMP* group increased by 44.93%. These findings suggest that *f*_AMP_ has no significant effect on the generation capacity of highquality AMP candidates. Merely optimizing antimicrobial activity is not sufficient to design AMPs with excellent properties. Compared to *M*_*A*_, the *High-quality* group in *M*_*AT*_ increased by 70.83%, the *Only-toxin* group increased by 34.58%, and the *Only-AMP* group decreased by 50.43%. These observations illustrate that optimizing both antimicrobial activity and cytotoxicity can significantly improve the quality of AMP candidates, and significantly reduce the AMP candidates that only satisfy partial property requirements. The same conclusion can also be drawn by comparing *M*_*AD*_ with *M*_*ATD*_. Through the comparison between *M*_*A*_ and *M*_*AD*_, as well as between *M*_*AT*_ and *M*_*ATD*_, we observe that *f*_DR_ has no significant impact on the composition of the four groups. These observations indicate that optimizing sequence diversity can significantly enhance the diversity of AMP candidates while not affecting the quality of the AMP candidates. In conclusion, extensive ablation studies highlight that specialized optimization of antimicrobial activity, cytotoxicity, and diversity can effectively enhance the performance of MPOGAN in generating high-quality AMP candidates.

### 2.7 Wet-laboratory validation

To further evaluate the MPOGAN-generated AMP candidates, we conducted wet-laboratory validation. The top ten high-confidence peptides screened out through the preliminary selection process (see “Methods”), were synthesized via solid-phase synthesis, and all purities are ≥95%. All ten peptides are novel and cannot be found in the existing UniProt [52] database (see Supplementary Table 4). We also synthesized a well-known AMP, LL37 [53], as a positive control; and a low-score peptide MPOP-Neg selected from MPOPs-50k, as a negative control. Sequence and structural information for these peptides can be found in Supplementary Table 3 and Supplementary Figs. 8 and 9. We tested the minimum inhibitory concentrations (MICs) of these 12 peptides against Gram-negative bacteria *E. coli* and two Gram-positive bacteria, *S. aureus* and *B. subtilis* (see “Methods”). A lower MIC value indicates a stronger bacterial inhibition effect. As shown in Fig. 5a, d, all synthesized peptides exhibit potent antimicrobial activities with MIC values of ≤512*µ*g/ml, while the MPOP-Neg shows a MIC value of ≥512*µ*g/ml. For ten high-confidence peptides, 80% exhibit MIC values against *B. aureus* of ≤16*µ*g/ml. All of these peptides show MIC values of ≤32*µ*g/ml against *S. aureus* and 80% of them against *E. coli*. Specifically, MPOP-03 and MPOP-04 showed the best antimicrobial activity among all synthesized peptides, with broad-spectrum antimicrobial effects over LL37 against all bacterial strains tested. These observations demonstrate the exceptional performance of MPOGAN in generating novel AMPs with broad-spectrum antimicrobial activity surpassing known AMPs.

**Fig. 5.**
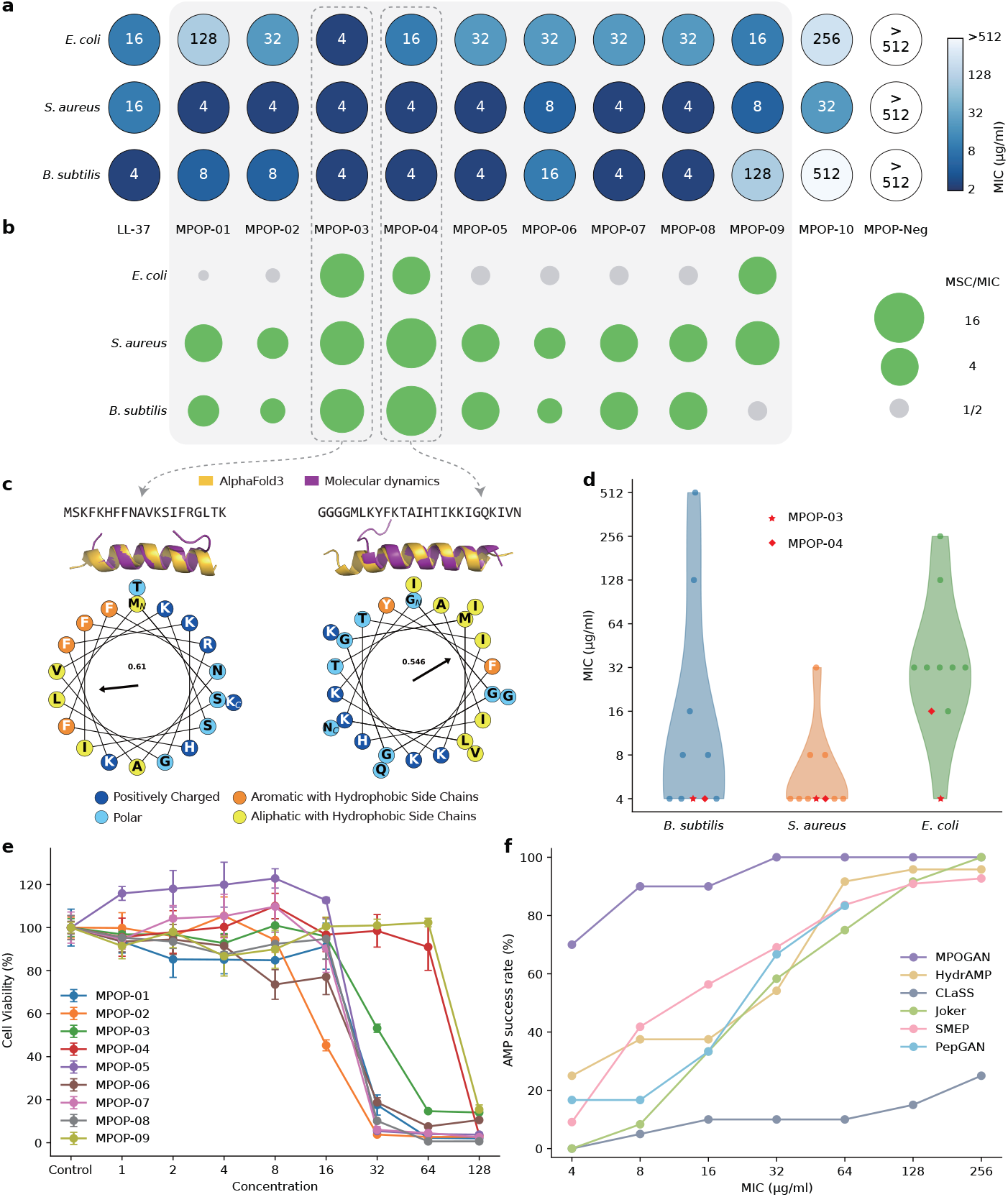
Wet-laboratory validation of the MPOGAN-generated AMPs. **a** MIC values of synthesized AMPs (x-axis) tested against pathogens including *E. coli, S. aureus*, and *B. subtilis* measured in duplicates (*n* = 3 independent experiments). **b** Safety assessment of bacterial infection treatment. MSC refers to the maximum safe concentration for eukaryotic cells. A larger MSC/MIC indicates a safer treatment. **c** 3D structures (created by AlphaFold 3 and molecular dynamics simulation) and alpha-helical wheels of two potent AMPs (MPOP-03 and MPOP-04). The numbers in the wheels represent Eisenberg hydrophobic moment. **d** Comparison of the antimicrobial effects of the MPOGAN-generated AMPs against pathogens in **a. e** Cell viability of MPOPs at various concentrations (*n* = 3 independent experiments, mean ±s.d.). The data were plotted as AMP concentration versus the percentage of living cells (see “Methods”). **f** Success rates in AMP wet-laboratory validation of MPOGAN, in comparison to HydrAMP, ClaSS, Joker, SMEP, and PepGAN at various MIC thresholds.

To evaluate the cytotoxicity of AMP candidates generated by MPOGAN, we tested the cytotoxicity of nine MPOPs (MPOP-01 to MPOP-09) against MC3T3-E1 cells. These nine peptides exhibit a better inhibitory effect than LL37 against some pathogens. As shown in Fig. 5e, we measured the cell viability (see “Methods”) of the nine peptides at various concentrations. As shown in Fig. 5b, we defined the maximum concentration value at which cell viability exceeded 50% as the maximum safe concentration (MSC) and calculated the MSC/MIC of the nine MPOPs against three bacterial strains. All nine MPOPs exhibit MSC/MIC≥2 against *S. aureus*, among which eight MPOPs exhibit MSC/MIC ≥1 against *B. subtilis*, suggesting that the peptides generated by MPOGAN have the potential to treat Gram-positive bacterial infections without damaging eukaryotic cells. Especially, MPOP-03 and MPOP-04 can effectively inhibit all tested pathogens, while avoiding damage to eukaryotic cells. The observations indicated that the nine MPOPs exhibit low cytotoxicity while effectively inhibiting bacterial pathogens, thus reducing eukaryotic cell damage and minimizing dosage requirements for local administration.

Furthermore, we predicted 3D structures of MPOP-03 and MPOP-04 using AlphaFold3 [54]. We then conducted 100ns molecular dynamics simulations (see Supplementary Note 4, and Supplementary Fig. 10) using Gromacs 2024.2 [55]. As shown in Fig. 5c top, both MPOP-03 and MPOP-04 exhibit stable alpha-helical structures. The alpha-helical wheels (Fig. 5c bottom) show strong amphiphilicity: positively charged amino acids (H, K, R) and polar amino acids (N, Q, S, T) aggregate to form the hydrophilic side, and hydrophobic amino acids (A, I, L, V, M, F, Y) aggregate to form the hydrophobic side. This amphiphilic distribution enables AMPs to selectively bind and disrupt bacterial cell membranes, enhancing their antimicrobial efficacy. More structural information of synthesized peptides can be found in Supplementary Figs. 8 and 9.

We also compared the success rates in AMP wet-laboratory verification under various MIC thresholds for MPOGAN and five state-of-the-art methods (HydrAMP [10], CLaSS [11], Joker [56], SMEP [4], and PepGAN [32]) (Fig. 5f). Statistics are obtained from the experimental data reported by these methods, regardless of differences in the target bacterial strains measured. MPOGAN outperforms other methods under all MIC thresholds. Under the MIC threshold of 32*µ*g/ml (potent antimicrobial activity), the success rate of MPOGAN has already reached 100%, whereas the success rates of other methods are all below 70%. Even under a MIC threshold of 4ug/ml (extremely potent antimicrobial activity), the success rate of MPOGAN remains strong at 70%, while all other methods struggle to exceed 25%. Although HydrAMP, CLaSS, and SMEP have success rates of ≥90% under 128 and 256 MIC thresholds (relatively broad conditions), these methods do not perform well under smaller MIC thresholds. In summary, the success rates in AMP wet-laboratory validation of MPOGAN are significantly better than other state-of-the-art methods.

## 3 Discussion

In this study, we develop and establish MPOGAN, a novel GAN framework for de novo design of AMPs with potent antimicrobial activity, reduced cytotoxicity, and diversity. The MPOGAN framework consists of two learning stages, including the pretraining stage and the multi-property optimizing (MPO) stage. The pre-training stage enables MPOGAN to learn the general characteristics of experimentally validated AMPs. In the MPO stage, we constructed a dynamic dataset for the adversarial finetuning of MPOGAN. We designed a model-embedded screening process that combines multiple plug-and-play attribute evaluators, providing high-quality generated data with multiple desired properties for the dynamic dataset. We also proposed a robust real-time knowledge-updating strategy to guide the updating of the dynamic dataset, thereby encouraging MPOGAN to iteratively fine-tune on new high-quality data while avoiding forgetting the old ones. Through the two learning stages, MPOGAN enhances the ability to generate AMPs that meet the antimicrobial activity, cytotoxicity, and diversity requirements. With these advancements, MPOGAN significantly outperforms baseline methods in generating AMPs de novo. Nine out of ten AMP candidates synthesized in the laboratory exhibit broad-spectrum antimicrobial activity and low cytotoxicity. Remarkably, the success rate of experimental validation of MPOGANdesigned AMPs exceeds other baseline methods, especially under severe restrictions.

## 4 Methods

### 4.1 Data collection

Our work’s training data comprises 30,812 peptide sequences, including an equal number of AMPs (15,406) and non-AMPs (15,406). AMPs are collected from known AMP databases, and non-AMPs are collected from the UniProt database [52]. To ensure both the availability of training data and the practicality of peptide synthesis in the laboratory, we limit the selected peptides to a maximum length of 25 amino acids following existing studies [7, 10].

For positive samples (i.e., real AMPs), we collect experimentally validated peptides from SATPdb [57], CAMP_R4_ [58], dbAMP [59], DRAMP [40], and LAMP [60] databases. We remove duplicate peptides and those ≥25 amino acids or containing non-standard amino acids. Negative samples (i.e., non-AMPs) are manually selected through the search criteria of UniProt [52]. These criteria necessitate the subcellular location to be the cytoplasm and exclude any features such as antimicrobial, antibiotic, antiviral, antifungal, effector, and excreted [10]. To increase the diversity of non-AMPs, negative samples with ≥40% sequence similarity are discarded and replaced with representative sequences from the cluster using CD-HIT [43]. Following the same method used in [10], the negative samples retain the same length distribution and size as the positive samples, thus reducing the bias between the two sample types.

Furthermore, to train and evaluate the antimicrobial activity predictor, we divide the dataset into training, validation, and test sets in an 8:1:1 ratio. Each subset maintains an equal balance of positive and negative samples. The amino acid composition and length distribution of the dataset are shown in Supplementary Fig. 1.

### 4.2 Antimicrobial activity predictor

To identify the highly potent AMP candidates, we construct an antimicrobial activity predictor, called LLM-AAP, based on ESM-2 [42]. As a ProLLM, ESM-2 has demonstrated excellent performance in various downstream tasks [7, 61, 62]. It leverages a BERT [63] encoder and is trained with 150 million parameters unsupervised on the Uniref50 [64] dataset. The LLM-AAP is modeled as a downstream task of the pretrained ESM-2 encoder network. We employ ESM-2 to encode peptides into feature representations, which are subsequently fed into a Multilayer Perceptron (MLP) to predict the probabilities of peptides being AMPs. In detail, the peptide sequence, denoted as *x* and with a length of *L* amino acids, is encoded by ESM-2 to generate residue-level representations with a dimension of (*L*+2, *E*). Subsequently, an elementwise average operation is performed across the length dimension to produce the global sequence representation with a dimension of (1, *E*). This global representation effectively captures the comprehensive features of the protein sequence. The sequence representations are further processed through three fully connected layers with rectified linear unit (ReLU) [65] activations and batch normalization. The input layer of MLP has the same number of neurons as the output dimension from ESM-2. The final layer of MLP uses a softmax function to enable class prediction. The LLM-AAP employs an Adam optimizer and trains on binary cross-entropy loss for 200 epochs. And the ESM-2 model parameters remain fixed throughout the training process.

We employ the optimal hyperparameters obtained from the grid search, featuring a learning rate of 5 ×10^−5^, a batch size of 256, and three hidden layers within the MLP, containing 256, 128, and 64 neurons, respectively. We set the dropout rate at a consistent 0.1.

### 4.3 Pre-training

MPOGAN is an extended GAN model comprised of a generator *G*_*θ*_ (Fig. 1b) and a discriminator *D*_*ϕ*_ (Fig. 1c), with model parameters represented as *θ* and *ϕ* respectively. Specifically, *G*_*θ*_ generates a sequence of amino acid tokens from a start token *s*_0_, while *D*_*ϕ*_ assesses whether the sequence is real. *G*_*θ*_ and *D*_*ϕ*_ engage in adversarial training until *G*_*θ*_ can generate sequences that are real enough that *D*_*ϕ*_ cannot distinguish whether they are real or generated sequences.

To enable MPOGAN to learn the features of real AMPs and generate peptides similar to real AMPs, we establish a pre-training stage that includes: generator pretraining, discriminator pre-training, and adversarial learning of *G*_*θ*_ and *D*_*ϕ*_.

#### 4.3.1 Generator pre-training

*G*_*θ*_ is a Recurrent Neural Network (RNN) with three layers. The first layer is a 3-dimensional embedding layer for each amino acid, leveraging word embeddings to capture semantic relationships among amino acids and reduce computational complexity. The next layer is a Gated Recurrent Unit (GRU) with 128 units. By introducing an update gate and a reset gate, the GRU retains historical information while controlling the impact of new inputs. Finally, the model ends with a fully connected layer using a softmax activation. This layer maps the GRU output to all possible amino acids, and softmax activation ensures these predictions sum to one, which can be interpreted as probabilities for each amino acid to be the next one in the sequence. We define the initial state of *G*_*θ*_ as *s*_0_ = 0.

We pre-train *G*_*θ*_ on the real AMP dataset, enabling it to have basic generative abilities. We also use maximum likelihood estimation to minimize the negative log-likelihood loss. The Adam optimizer is employed to update the model parameters, with an initial learning rate of 0.0001. *G*_*θ*_ is trained for a total of 1000 epochs.

#### 4.3.2 Discriminator pre-training

*D*_*ϕ*_ functions as a binary classifier that distinguishes between real and generated sequences. Instead of additionally pre-training a *D*_*ϕ*_ separately, we directly employ the parameters of the trained LLM-AAP as *ϕ*.

#### 4.3.3 Adversarial learning

To generate peptides that can deceive *D*_*ϕ*_, *G*_*θ*_ acts as an actor and outputs a complete peptide sequence *x* starting from *s*_0_ to maximize the expected cumulative reward. To avoid the differentiation difficulty associated with discrete data in conventional GAN, the *G*_*θ*_ is trained via policy gradient [41]. We calculate the loss of *G*_*θ*_ as follows:

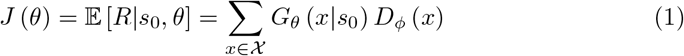

where *R* stands for the reward of a complete AMP sequence, and χ stands for the entire sequence space. The reward function *R* is defined as the positive probability output of *D*_*ϕ*_ for the generated sequence.

The optimization objective of *D*_*ϕ*_ is to maximize the probability of accurately classifying both real and generated peptides. We calculate the loss of *D*_*ϕ*_ as follows:

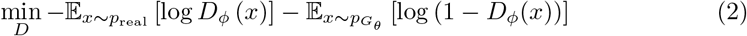

where *p*_real_ stands for the distribution of real AMPs, and 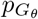 stands for the distribution of *G*_*θ*_. In our work, the adversarial learning process is conducted for 1000 epochs.

### 4.4 Multi-property optimizing

To further enhance the ability of the pre-trained MPOGAN to generate AMP candidates with multiple desired properties, we establish a multi-property optimizing (MPO) stage, including model-embedded screening, Real-Time Knowledge-Updating (RTKU), and adversarial fine-tuning. Model-embedded screening is proposed to identify high-quality peptides that meet the multiple desired properties from generated AMP candidates. The RTKU strategy implements an extended feedback-loop mechanism, which combines AMP candidates screened from model-embedded screening and a random set of real AMPs. It further updates these two sets into the dynamic dataset and adjusts the size of the AMP candidates set fed into the model-embedded screening in the next iteration. Adversarial fine-tuning is designed to enable MPOGAN to iteratively learn the features of high-quality AMP candidates on dynamic datasets, improving the performance of generating multiple desired properties.

#### 4.4.1 Model-embedded screening

The prerequisite for fine-tuning MPOGAN on the dynamic dataset is to ensure data quality. In our work, we introduce model-embedded screening to address this challenge. When the *τ* th iteration begins, a total of *N*_*τ*_ peptides (*x*) are sampled from *G*_*θ*_.

We abstract these peptides as a set 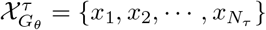. Deep generative models are highly susceptible to the impact of noise interference [36]. Therefore, the peptides utilized to update the dynamic dataset should be of high confidence to avoid unexpected bias. To screen out high-quality AMP candidates from 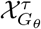, we consider the antimicrobial activity, cytotoxicity, and diversity as the key properties to be optimized.

Firstly, we employ the antimicrobial activity predictor (*f*_AMP_) to screen the antimicrobial activity of 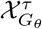, given its fundamental importance for AMPs. Candidates with a probability of antimicrobial activity *>*0.8 are retained.

Secondly, the avoidance of toxic peptides, which poses a significant challenge in AMP research [66], necessitates further prediction of cytotoxicity for qualifying peptides. We employ ToxinPred2 [18] (see Supplementary Note 1.1) as the cytotoxicity predictor (*f*_Toxin_). Candidates with a probability of cytotoxicity *<*0.7 are retained.

Thirdly, to remove redundancy from candidates and enrich the diversity of the dynamic dataset, we utilize CD-HIT [43] (see Supplementary Note 1.2) as the de-redundancy module, represented as the de-redundancy function *f*_DR_. Representative candidates with sequence similarity *<*60% are retained.

Finally, we define the model-embedded screening as follows:

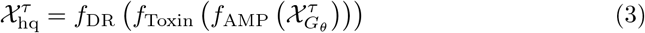

where 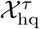 stands for the high-quality AMP candidates outputted from model-embedded screening in the *τ* th iteration.

#### 4.4. 1 Real-Time Knowledge-Updating strategy

In the *τ* th iteration of the MPO stage, we define the dynamic dataset as χ^*τ*^, and the peptides to be updated as 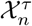.To ensure the robustness of the update strategy, we fix the size of the dynamic dataset to 1000 (|χ^*τ*^| = 1000, *τ* ≥ 0), and the size of the peptides to be updated to 250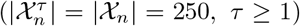. Specifically, χ^0^ is randomly initialized with the real AMP dataset (χ_real_). The RTKU strategy for χ^*τ*^ adheres to the first-in-first-out (FIFO) principle:

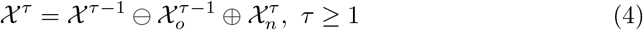

where ⊖ stands for the set difference operation, ⊕ stands for the set union operation, and 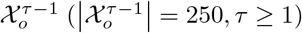 stands for the oldest updated peptides in χ^*τ*−1^.

More details of the update strategy of χ^*τ*^ can be found in Supplementary Note 1.3 and Supplementary Fig. 2.

Similar to incremental learning (IL) [67], the original feedback-loop mechanism continuously extracts knowledge from the new data generated by GAN. However, this paradigm has caused two problems: (1) Mode collapse. The homogenization of peptides updated into the dynamic dataset makes it difficult for GAN to learn diverse knowledge and generate diverse peptides. (2) Catastrophic forgetting. Directly optimizing the network with new knowledge will erase the knowledge from earlier stages and result in irreversible performance degradation [68]. In this study, we address the mode collapse problem by introducing a de-redundancy module in the model-embedded screening. To address the catastrophic forgetting problem, the RTKU strategy fine-tunes MPOGAN using both new data from peptides generated by MPOGAN and old data from χ_real_, inspired by replay-based IL [67]. Specifically, we introduce a hyperparameter *α* = 0.7 to balance the relative size of new data 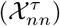 and old data 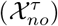 in 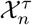:

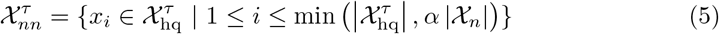

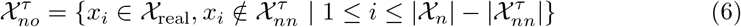

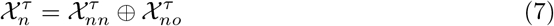

To ensure the robustness and efficiency of the RTKU strategy, *N*_*τ*_ is designed as a value that varies with iteration to ensure that model-embedded screening outputs a sufficient amount of new knowledge while minimizing screening time as follows:

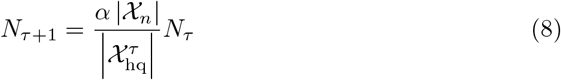

#### 4.4.3 Adversarial fine-tuning

The adversarial fine-tuning of MPOGAN is performed on the dynamic dataset χ^*τ*^, with all sequences in the dataset considered positive samples. The training of *G*_*θ*_ and *D*_*ϕ*_ is consistent with the pre-training stage. *G*_*θ*_ is trained to generate high-quality AMP candidates, while *D*_*ϕ*_ is trained to distinguish AMPs and non-AMPs.

Through repeated iterations of model-embedded screening and adversarial finetuning, MPOGAN continuously learns the features of AMP candidates with potent antimicrobial activity, low cytotoxicity, and high diversity simultaneously. On the one hand, *D*_*ϕ*_ learns the features of high-quality AMP candidates, which leads to a stricter definition of “positive samples” and continuous updates to the parameters *ϕ*, resulting in more precise discrimination of the sequences generated by *G*_*θ*_. On the other hand, *G*_*θ*_ continuously updates the parameters *θ* based on the feedback from *D*_*ϕ*_, optimizing the generation preferences to generate peptides that are more similar to the dynamic dataset, i.e., peptides that possess high-quality features.

### 4.5 Wet-laboratory validation

#### 4.5.1 Preliminary selection of potential candidates for Wet-laboratory validation

Although most of the AMPs directly generated by MPOGAN exhibit potent antimicrobial activity and reduced cytotoxicity, we still established a preliminary selection process (Supplementary Fig. 7). This process aims to select the most promising AMP candidates generated by MPOGAN for wet-laboratory validation.

For the AMP candidates generated by MPOGAN, we first remove the sequences that already exist in the real AMP dataset. To preliminarily exclude sequences that do not conform to expert experience, we perform the following biological experience screening [4, 10]: (1) screening AMP candidates with a positive net charge; (2) excluding AMP candidates in which there are more than three positively charged amino acids (K, R) in a window of five amino acids; (3) excluding AMP candidates in which occur three hydrophobic amino acids in a row. We consider amino acids as hydrophobic based on the Eisenberg scale: A, V, L, I, M, F, W. To further ensure the key properties of AMP candidates, we introduce the auxiliary classifier screening. In detail, to ensure that the AMP candidates have high antimicrobial activity, we filter the AMP candidates using multiple antimicrobial activity predictors, including our antimicrobial activity predictor, AMPScanner v2 [16] and AMPlify [17]. We retain sequences with a probability of being an AMP≥ 0.99 predicted by all three predictors. To ensure that the AMP candidates have low cytotoxicity, we filter the sequences using the cytotoxicity predictor ToxinPred2 [18], retaining sequences with a cytotoxicity probability ≤ 0.5. To obtain sequences with an *α*-helix structure, which is a common structure of AMPs, we filter the sequences using the HeliQuest [69] web server. The parameters are set as follows: Hydrophobicity: -1 ≤ *H* ≤ 0.6; Hydrophobic moment: 0.4 ≤ *M* ≤ 1.2; Net charge: 0≤ *Ch* ≤ 5; Minimal number of polar residues + glycine: 0; Minimal number of uncharged residue: 1; Minimal number of glycine: 0; Maximal number of charged residue: 12; Is proline accepted? (at i, i+3 / n-3, n): Yes; Is cysteine accepted?: Yes.

Finally, to select the most promising AMP candidates for wet-laboratory validation, we calculate the weighted score for each sequence (*x*). The weighted score is calculated as follows:

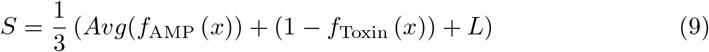

where *S* stands for the weighted score of the sequence, *Avg* (*f*_AMP_ (*x*)) stands for the average score of the three antimicrobial activity predictors (LLM-AAP, AMP Scanner v2, AMPlify), *f*_Toxin_ (*x*) stands for the score of the cytotoxicity predictor, and *L* stands for the length evaluation score. *L* is calculated as follows:

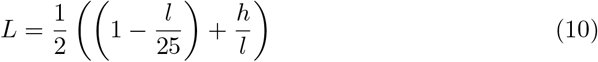

where *l* stands for the length of *x*, and *h* stands for the length of the helix in *x*. For the 50,000 AMP candidates generated from MPOGAN, the number of candidates is reduced to 49,538 after de-duplication. Then, with biological experience screening [4, 10], the candidate pool is further decreased to 18,153. Next, auxiliary classifier screening is conducted, and the number of candidates declines to 429. After further evaluating the secondary structure, we leave 124 peptides as stringent AMP candidates for physicochemical features analysis. Finally, we select the top 10 candidates with the highest weighted score for wet-laboratory validation.

#### 4.5.2 Antimicrobial activity assay

The antimicrobial activity of the synthesized AMPs is tested on strains of *E. coli* (BL21(DE3)), *S. aureus* (BNCC186335), and *B*.*subtilis* (BNCC109047). All strains are stored at -80°C and, before testing, are transferred into fresh Mueller-Hinton Medium and incubated for 24 h at 37°C. Fresh cultures are used for the antimicrobial assays. The MIC values are based on the broth microdilution method according to the Clinical and Laboratory Standards Institute (CLSI) protocol. Initial bacterial inoculums (0.510^5^ colony forming units (CFU)/mL) in Mueller-Hinton Broth are exposed to varying concentrations of the compounds (0.5-512 *µ*g/mL) and incubated for 18 h at 37°C. The experiments are conducted on 96-well microtiter plates with a final volume of 100 *µ*L. The MIC is defined as the lowest concentration of the compound that inhibits visible bacterial growth. Additionally, the MBC is evaluated. After determining the MIC, samples from the MIC well and two higher concentrations are plated on agar plates. The MBC is recorded as the lowest concentration at which no visible bacterial colonies are observed. All experiments are performed in triplicate, with positive controls (ensuring proper bacterial growth) and negative controls (ensuring sterility) included.

#### 4.5.3 Cytotoxicity assay

For the cytotoxicity test, 2,000 MC3T3-E1 cells were seeded into each well of 96-well plates and cultured in *α*-MEM medium containing 10% FBS and 1% penicillin-streptomycin at 37^*°*^C and 5% CO_2_ overnight. Then the culture medium was removed and the AMPs at the various concentrations in the media were added to each well. The cells were further incubated for 48 h, followed by the addition of 20 *µ*L of CCK-8 and subsequent incubation for 1 h. The absorbance of the solution was measured at 450 nm by a microplate reader. The cell viability was calculated as follows:

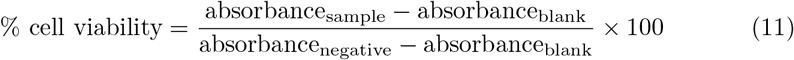

## 5 Data availability

All relevant datasets in this article are publicly available. SATPdb is available at http://crdd.osdd.net/raghava/satpdb/. CAMP_R4_ is available at https://camp.bicnirrh.res.in/. dbAMP is available at https://awi.cuhk.edu.cn/dbAMP/. DRAMP is available at http://dramp.cpu-bioinfor.org/. LAMP is available at http://biotechlab.fudan.edu.cn/database/lamp/. UniProt is available at https://www.uniprot.org/.

## 6 Code availability

The code is available on GitHub (https://github.com/23AIBox/MPOGAN).

