## SupplementaryInformation for "A Multi-Property Optimizing Generative Adversarial Network for de novo Antimicrobial Peptide Design"

### Supplementary Note

#### 1 Supplementary Methods

##### 1.1 Details of the cytotoxicity predictor

In our work, we utilized a hemolytic toxicity predictor to evaluate the cytotoxicity of peptides during the multi-property optimization (MPO) process, guiding MPOGAN to learn the features of peptides with low cytotoxicity. We employ the trained ToxinPred2 [1] as our cytotoxicity predictor. ToxinPred2 is an enhanced method designed to predict the toxicity of peptides and small proteins, advancing on the original ToxinPred [2]. ToxinPred2 employs techniques including Basic Local Alignment Search Tool-based similarity, Motif-EmeRging and with Classes-Identification-based motif search, along with prediction models. By combining these strategies, ToxinPred2 achieves high accuracy, maintaining a balance between sensitivity and specificity.

##### 1.2 Details of the sequence de-redundancy module

During the model-embedded screening process, it is crucial to constantly update the dynamic dataset in real-time, enabling MPOGAN to learn the features of high-quality peptides. However, if a peptide has excellent properties, it is likely that peptides with similar sequences have similar properties. This can result in a higher similarity between high-quality peptides identified by both the antimicrobial activity predictor and the cytotoxicity predictor during the model-embedded screening process. If we employ peptides screened by the initial two screening modules directly to update the dynamic dataset, it may diminish the diversity of high-quality peptides, leading to a homogenization of features learned by MPOGAN. To address this issue, we incorporate a sequence de-redundancy module as the final component of the model-embedded screening process. Given its impact on screening efficiency, we utilize CD-HIT [3] as the sequence de-redundancy module.

CD-HIT, an advanced program for biological sequence clustering, significantly speeds up the clustering process through a unique parallelization strategy and other enhancements. By streamlining core steps, implementing comprehensive parallelization with multiple threads, and integrating improvements like quicker file reading and advanced filtering, CD-HIT achieves notable acceleration. This enables efficient clustering of vast datasets in a fraction of the time compared to earlier versions.

##### 1.3 More information on the Real-Time Knowledge-Updating (RTKU) strategy

The dynamic dataset is updated with iterations of the MPO stage. In our work, as shown in Supplementary Fig. 1, the dynamic dataset consists of 1000 peptides, with the oldest 250 peptides being updated in each iteration. We divided the dataset into 4 groups of equal size, each containing 250 peptides. With this grouping, only the currently oldest sequence needs to be substituted with the latest peptides in each iteration. In the dynamic dataset, the entry time of each peptide is recorded and updated in sync with the dataset. Before the MPO stage, the dynamic dataset is initialized with 1000 randomly selected data from the real AMPs dataset, and the entry time of all the peptides is initialized to 0. Beginning from the first iteration, the oldest 250 peptides are removed in each round based on a first-in-first-out principle, and subsequently replaced with 250 new peptides. The entry time of the newly added peptides is set to the current iteration. The entire dynamic dataset will be completely updated once every 4 iterations.

#### 2 Baseline methods and evaluation settings

##### 2.1 Baseline methods for AMPs generation

In the AMPs generation task, we compare five baseline methods, which are Dean-VAE [4], Nagajaran-LSTM [5], PepCVAE [6], HydrAMP [7], and Muller-RNN [8].

**Dean-VAE.** Dean-VAE trains a variational autoencoder (VAE) on a database of known antimicrobial peptides (AMPs) and their scrambled counterparts to generate a continuous latent space representation of AMPs. The encoder network converts peptide sequences into latent space vectors and the decoder network converts these vectors back into peptide sequences. New AMP sequences are generated by sampling points in the latent space near known active AMPs. The model's capability to generate novel active AMPs is validated by testing the antimicrobial activity of interpolated sequences between known active and inactive peptides.

**Nagajaran-LSTM.** Nagajaran et al. implement a computational approach that employs a long short-term memory (LSTM) language model to design antimicrobial peptides. The LSTM model is applied to analyze the arrangement and frequencies of amino acid residues in known antimicrobial peptide sequences, interpreting them as words of a 20-alphabet language. Based on the output of the LSTM network, ten of LSTM-generated peptides are synthesized and tested against bacterial pathogens. The study demonstrates the effectiveness of this LSTM-based peptide design approach in generating peptides with broad-spectrum antimicrobial activity, including against multidrug-resistant clinical isolates of bacteria.

**PepCVAE.** PepCVAE is a semi-supervised VAE. This framework aims to design novel AMPs sequences by learning a rich latent space of biological peptide context from unlabeled peptide sequences. By leveraging feedback from a jointly trained AMPs classifier using limited labeled instances, the model further learns a disentangled antimicrobial attribute space, allowing for controllable generation of AMPs. The PepCVAE architecture demonstrates superior performance compared to a plain VAE by generating novel AMPs with higher long-range diversity while remaining closer to the training distribution of biological peptides.

Since the code, trained model, training data, and generated peptides for this work are not available, and Szymczak et al. [7] reproduced it successfully, the results reproduced by Szymczak et al. are used for comparison in our work.

**HydrAMP.** HydrAMP is a conditional variational autoencoder (cVAE) designed for AMPs generation. This model learns a lower-dimensional, continuous representation of peptides, disentangling their antimicrobial properties. It can generate analogues of existing peptides with specified antimicrobial conditions and also generate peptides de novo. HydrAMP leverages a continuous peptide representation with disentangled antimicrobial conditions, allowing for both analogue and unconstrained generation. The model benefits from a temperature parameter controlling creativity in analogue generation and has been trained specifically for the task of analogue generation, including generating both positive and negative analogues. In addition, the model introduces a preselection procedure based on external classifiers and molecular dynamics simulations, enhancing the experimental validation rate of the generated peptides.

**Muller-RNN.** Muller et al. developed a generative LSTM recurrent neural network (RNN) based on amino acid sequences of helical AMPs. The trained model was used for de novo sequence generation, resulting in 82% of the generated sequences predicted to be active antimicrobial peptides. Furthermore, the generated sequences were observed to be more similar to the training data than manually designed amphipathic helices. This showcases the potential of LSTM in constructing new amino acid sequences for peptide and protein design without the need for exhaustive enumeration of sequence libraries.

**Evaluation settings.** For Dean-VAE, Nagajaran-LSTM, and Muller-RNN, we collected the generated AMP candidates provided in their Supplementary Materials. For HydrAMP and PepCVAE, we used Supplementary Algorithm six of [7] to obtain their AMP candidates. Our evaluation for MPOGAN does not involve any screening processes but directly assesses the model-generated peptides. For MPOGAN, we sampled 50,000 peptides directly from the pre-trained MPOGAN and MPOGAN, respectively. To ensure a fair and consistent comparison, we excluded peptides longer than 25 amino acids and those containing non-standard amino acids. The final number of AMP candidates we collected for comparison is as follows: 2973 for Dean-VAE, 25,607 for Nagajaran-LSTM, 1052 for Muller-RNN, 50,000 for HydrAMP, 50,000 for PepCVAE, and 50,000 for MPOGAN.

In this evaluation, we apply two predictive models to evaluate the antimicrobial activity and toxicity of AMP candidates. Specifically, we employ our trained antimicrobial activity predictor (LLM-AAP) to assess the probability that AMP candidates have antimicrobial activity ($P_{AMP} \in [0,1]$). A larger $P_{AMP}$ value indicates a higher probability that an AMP candidate has antimicrobial activity. $P_{AMP} > 0.8$ is considered to have a high probability of having antimicrobial activity. Toxinpred2 [1] is used to assess the probability that AMP candidates have peptide toxicity ($P_{Toxin} \in [0,1]$). A smaller $P_{Toxin}$ value indicates a lower probability that an AMP candidate has peptide toxicity. $P_{Toxin} < 0.7$ is considered to have a low probability of having peptide toxicity. We expect to obtain AMP candidates that exhibit both high antimicrobial activity and low peptide toxicity, so the AMP candidates that meet both $P_{AMP} > 0.8$ and $P_{Toxin} < 0.7$ are considered high-quality AMP candidates. In addition, uniqueness is derived by counting the proportion of only one occurrence in the set of generated sequences.

##### 2.2 Baseline methods for AMPs identification

In the AMPs identification task, we compare six baseline methods, which are MACREL [9], amPEPpy [10], Ma-NNMs [11], AMPlify [12], STM [13], and AMP Scanner v2 [14].

**MACREL.** MACREL is a pipeline for predicting AMPs from genomes and metagenomes, utilizing two feature-based classifiers to predict AMP and hemolytic activity based on 22 descriptors. The classifiers incorporate a combination of local and global features, including amino acid distribution patterns, physiochemical properties, solubility, and composition of amino acid groups. Training sets for the classifiers are carefully curated to ensure accuracy, with AMP prediction using a random forest classifier and hemolytic activity prediction based on the HemoPI-1 dataset. MACREL processes metagenomic reads, assembles contigs, predicts genes, and classifies AMP sequences based on their characteristics. The pipeline demonstrates high precision and sensitivity in identifying high-quality AMP candidates, making it a valuable tool for extracting AMPs from genomic and metagenomic data.

**amPEPpy.** amPEP [15] is a computational method that predicts AMPs using the random forest algorithm. The method utilizes distribution patterns of amino acid properties along the sequence to develop a prediction model. A large and diverse dataset of AMP and non-AMP sequences is used to evaluate different random forest classifiers with varying positive:negative data ratios. The optimal model, amPEP, achieves high accuracy, Matthew's correlation coefficient (MCC), area under the receiver operating characteristic curve (AUROC), and the Kappa statistic. Feature analysis is conducted to identify a minimal set of 23 features for AMP prediction with high accuracy. The method outperforms existing approaches in terms of accuracy, MCC, and AUROC when tested on benchmark datasets.

In our paper, we compare model performance using amPEPpy, which is the Python 3 implementation of amPEP. amPEPpy has better portability and provides a command-line user interface designed to efficiently process genome-scale data.

**Ma-NNMs.** Ma et al. integrate multiple natural language processing neural network models, including LSTM, Attention, and BERT, to establish a unified pipeline for identifying candidate AMPs from human gut microbiome data. By treating peptide sequences as text data and leveraging large datasets, they use these neural network models (NNMs) to predict AMPs effectively. They optimize predictive performance by combining the strengths of each model and achieve high precision and recall rates in distinguishing AMPs from non-AMPs. Additionally, they mine metagenomic and metaproteomic data to filter and select potential AMP candidates for further validation, demonstrating the power of machine learning approaches in discovering functional peptides from metagenome data.

**AMPlify.** AMPlify is an attentive deep learning model for the prediction of AMPs. AMPlify utilizes a bidirectional long short-term memory (Bi-LSTM) layer to encode positional information, followed by a multi-head scaled dot-product attention layer for refined sequence representation. A context attention layer generates a summary vector by learning contextual information. The model is trained on known AMPs and non-AMPs, incorporating ensemble learning to enhance performance. Additionally, attention mechanisms are applied to improve in silico AMP prediction, with a focus on discovering novel peptide-based alternatives to conventional antibiotics.

**STM.** The "Sense the Moment" (STM) prediction system aims to differentiate between AMPs and shuffled versions by using hydrophobic moment values. Unlike deep learning methods that require large amounts of training data, STM focuses on the distinctive features of AMPs and shuffled peptides. By calculating the geometric average of hydrophobic moments measured on different scales, the system demonstrates high accuracy and sensitivity in predicting AMPs. This approach underscores the potential of utilizing specific physicochemical properties, such as hydrophobic moment, as effective classifiers in antimicrobial peptide prediction, presenting a promising alternative to traditional machine learning algorithms.

**AMP Scanner v2.** The Antimicrobial Peptide Scanner v2 (AMP Scanner v2) utilizes a deep neural network (DNN) model with convolutional and recurrent layers to identify antimicrobial peptides (AMPs) based on primary sequence composition. The DNN model processes peptide sequences that are encoded into uniform numerical vectors, which are then fed through an embedding layer, convolutional layer, and LSTM layer to capture position-invariant patterns and automatically extract features. This model's architecture enables efficient recognition of AMPs without requiring domain experts to construct features beforehand. Moreover, the model learns a reduced alphabet representation of peptide sequences to further improve the accuracy of AMP recognition.

**Evaluation settings.** The calculation of evaluation metrics for the AMP identification methods is as follows:

$$\text{Accuracy} = \frac{TP + TN}{TP + TN + FP + FN}$$

$$\text{Sensitivity} = \frac{TP}{TP + FN}$$

$$\text{F1 score} = \frac{2 \times \text{Precision} \times \text{Recall}}{\text{Precision} + \text{Recall}}$$

$$\text{MCC} = \frac{TP \times TN - FP \times FN}{\sqrt{(TP + FP)(TP + FN)(TN + FP)(TN + FN)}}$$

where $TP$, $TN$, $FP$, and $FN$ stand for true positive, true negative, false positive, and false negative, respectively. Precision is defined as $\frac{TP}{TP+FP}$, and Recall is defined as $\frac{TP}{TP+FN}$. The predictor is trained on the training set and evaluated on the validation set. The final performance is reported on the test set.

#### 3 Robustness assessment of MPO stage

During the MPO stage, the dynamic dataset is continuously updated. At each update, the newly added data changes the composition of the dataset before the update. We aim to avoid undesired biases and ensure that the model still learns the features of real AMPs. As shown in Supplementary Figure 2, we evaluated the changes in the composition of the dynamic training dataset during the MPO stage. The dynamic training dataset is initialized with real AMPs before the training starts, and the number of generated sequences and real AMPs change rapidly in the first 4 iterations, and remain stable in the subsequent iterations, with slight fluctuations between 650-700 and 300-350, respectively. This proves that the MPO process is robust, and the dataset is able to maintain a stable compositional state while being dynamically updated.

#### 4 Molecular dynamic simulations

Our molecular dynamics (MD) simulation began with the initial structure obtained through peptide structure prediction using AlphaFold3 [16]. This simulation was executed utilizing Gromacs 2024.2 [17] software. The simulation box was filled with TIP3P water [18], and NaCl ions were introduced to neutralize the system. Steric clashes were eliminated through energy minimization using the steepest descent algorithm. The simulations were conducted using a time step of 2 femtoseconds, ensuring accuracy in the integration of Newton's equations of motion. The LINCS algorithm [19] was used to constrain bonds involving hydrogen atoms. Electrostatic interactions were calculated using the Particle-Mesh Ewald (PME) algorithm [20], with a real-space cut-off for pairs more than 1.2 nm apart. Lennard-Jones interactions were gradually switched to zero between 1 and 1.2 nm using the force-switch algorithm [21]. During the equilibration stages, pressure coupling was managed by the Berendsen barostat [22], which was later transitioned to the Parrinello-Rahman barostat [23] for all production stages. The barostat time constant and compressibility factor were uniformly set to 5 ps and $4.5 \times 10^{-5}$ $bar^{-1}$, respectively. The v-rescale thermostat [24] maintained the temperature at 310.15 K. After a 100ns simulation period, the root mean square deviation (RMSD) was computed to evaluate the stability of simulations. The MD simulation results were then visualized using PyMOL.

### Supplementary Tables

**Supplementary Table 1** Performance evaluation of methods for predicting antimicrobial peptides (AMPs) on the independent test set. Bolded numbers are the best performance.

| Methods | Accuracy (%) | Precision (%) | F1 score (%) | Sensitivity (%) | Specificity (%) | AUC | MCC (%) |
|---|---|---|---|---|---|---|---|
| STM | 57.79 | 56.92 | 60.27 | 64.05 | 51.52 | 0.5779 | 15.70 |
| MACREL | 76.83 | 93.39 | 71.37 | 57.75 | 95.91 | 0.8541 | 58.06 |
| amPEPpy | 77.06 | 80.84 | 75.56 | 70.93 | 83.19 | 0.8253 | 54.53 |
| Ma-NNMs | 78.49 | **94.08** | 73.87 | 60.80 | **96.17** | 0.7849 | 60.91 |
| AMPlify | 80.47 | 83.90 | 79.43 | 75.41 | 85.53 | 0.8695 | 61.25 |
| AMP Scanner v2 | 84.00 | 87.64 | 83.19 | 79.17 | 88.84 | 0.9082 | 68.33 |
| LLM-AAP (Ours) | **88.87** | 91.54 | **88.50** | **85.66** | 92.08 | **0.9517** | **77.90** |

**Supplementary Table 2** Percentage of AMP candidates generated by pre-trained MPOGAN (top row) and MPOGAN (bottom row) in different categories of Expect value (E-value). The E-value for the match with the highest score was considered, as obtained by performing a BLAST similarity search against the real AMPs dataset. Each model generates 50,000 AMP candidates for validation.

| E-value | <=0.0001 | <=0.001 | <=0.01 | <=0.1 | <=1 | <=10 | >10 |
|---|---|---|---|---|---|---|---|
| SeqGAN | 3.41 | 1.40 | 3.16 | 5.43 | 13.14 | 35.16 | 38.30 |
| MPO-GAN | 2.83 | 3.22 | 6.16 | 11.70 | 25.17 | 36.16 | 14.77 |

**Supplementary Table 3** Sequence information and physicochemical properties of synthesized peptides.

| Name | Sequence | length | Charge | Isoelectric point | Aromaticity | Eisenberg hydrophobicity | Hydrophobic moment | Hydrophobic ratio | Charge density | Instability index | Aliphatic index |
|---|---|---|---|---|---|---|---|---|---|---|---|
| LL-37 | LLGDFFRKSKEKIGKEFKRIVQRIKDFLRNLVPRTES | 37 | 7 | 11.1514 | 0.1081 | 5.7999 | 0.5624 | 0.3514 | 0.0013 | 23.3432 | 89.4595 |
| MPOP-01 | DPFGIMSKLQQFIRKFYQSLKHLKT | 25 | 5.094 | 10.7490 | 0.16 | -0.0624 | 0.4582 | 0.36 | 0.0013 | 69.6404 | 78 |
| MPOP-02 | LHQIKSVIKTAMNVLSGLFSAIKKK | 25 | 6.094 | 11.2788 | 0.04 | 0.1256 | 0.4815 | 0.48 | 0.0018 | -0.588 | 124.8 |
| MPOP-03 | MSKFKHFFNAVKSIFRGLTK | 20 | 6.094 | 11.7510 | 0.2 | 0.0025 | 0.6101 | 0.45 | 0.0021 | 13.085 | 58.5 |
| MPOP-04 | GGGGMLKYFKTAIHTIKKIGQKIVN | 25 | 6.093 | 10.9063 | 0.08 | 0.1316 | 0.5456 | 0.36 | 0.0019 | 3.336 | 93.6 |
| MPOP-05 | VMKRIGTILSGLHSLLSKIF | 20 | 4.094 | 11.5723 | 0.05 | 0.3005 | 0.5557 | 0.5 | 0.0014 | 26.94 | 151 |
| MPOP-06 | KLFRVVKKMFHSVFSSIHKYFR | 22 | 7.192 | 11.5869 | 0.2273 | -0.0382 | 0.7441 | 0.4545 | 0.0022 | 44.3227 | 75 |
| MPOP-07 | GIGKFLHSFTKFFSKIMNAIR | 21 | 5.094 | 11.6768 | 0.1905 | 0.1776 | 0.6721 | 0.4762 | 0.0017 | -2.6048 | 79.0476 |
| MPOP-08 | GLFDVIKKIAEMVSNGYHTVKKKF | 24 | 4.095 | 10.2998 | 0.125 | 0.0638 | 0.5509 | 0.4167 | 0.0011 | -1.35 | 89.1666 |
| MPOP-09 | IKGIIKTIKKMASHFLHSAS | 20 | 5.193 | 11.1567 | 0.05 | 0.1435 | 0.5760 | 0.45 | 0.0019 | 5.56 | 107.5 |
| MPOP-10 | IGSFKHVFKRITSMAKAIR | 19 | 6.094 | 12.1797 | 0.1053 | -0.0132 | 0.4946 | 0.4737 | 0.0023 | 17.3053 | 87.3684 |
| MPOP-Neg | LARGDCIMKLL | 11 | 1.929 | 8.7432 | 0 | 0.1664 | 0.3288 | 0.6364 | 0.0008 | -6.3455 | 150.9091 |

**Supplementary Table 4** BLAST sequence similarity search of synthesized peptides in the UniProt database.

| Peptide | Min E-value |
|---|---|
| MPOP-01 | 0.059 |
| MPOP-02 | 0.62 |
| MPOP-03 | 0.16 |
| MPOP-04 | 3.4 |
| MPOP-05 | 5.0 |
| MPOP-06 | 0.84 |
| MPOP-07 | 2.5 |
| MPOP-08 | 0.24 |
| MPOP-09 | 3.4 |
| MPOP-10 | 1.1 |

E-value < $1 \times 10^{-100}$: Identical sequences.
$1 \times 10^{-100}$ < E-value < $1 \times 10^{-50}$: Almost identical sequences.
$1 \times 10^{-50}$ < E-value < $1 \times 10^{-10}$: Closely related sequences, could be a domain match or similar.
$1 \times 10^{-10}$ < E-value < $1$: Could be a true homologue but it is a gray area.
$1$ < E-value < $10$: Proteins are most likely not related
E-value > $10$: Hits are most likely junk unless the query sequence is very short.

### Supplementary Figures

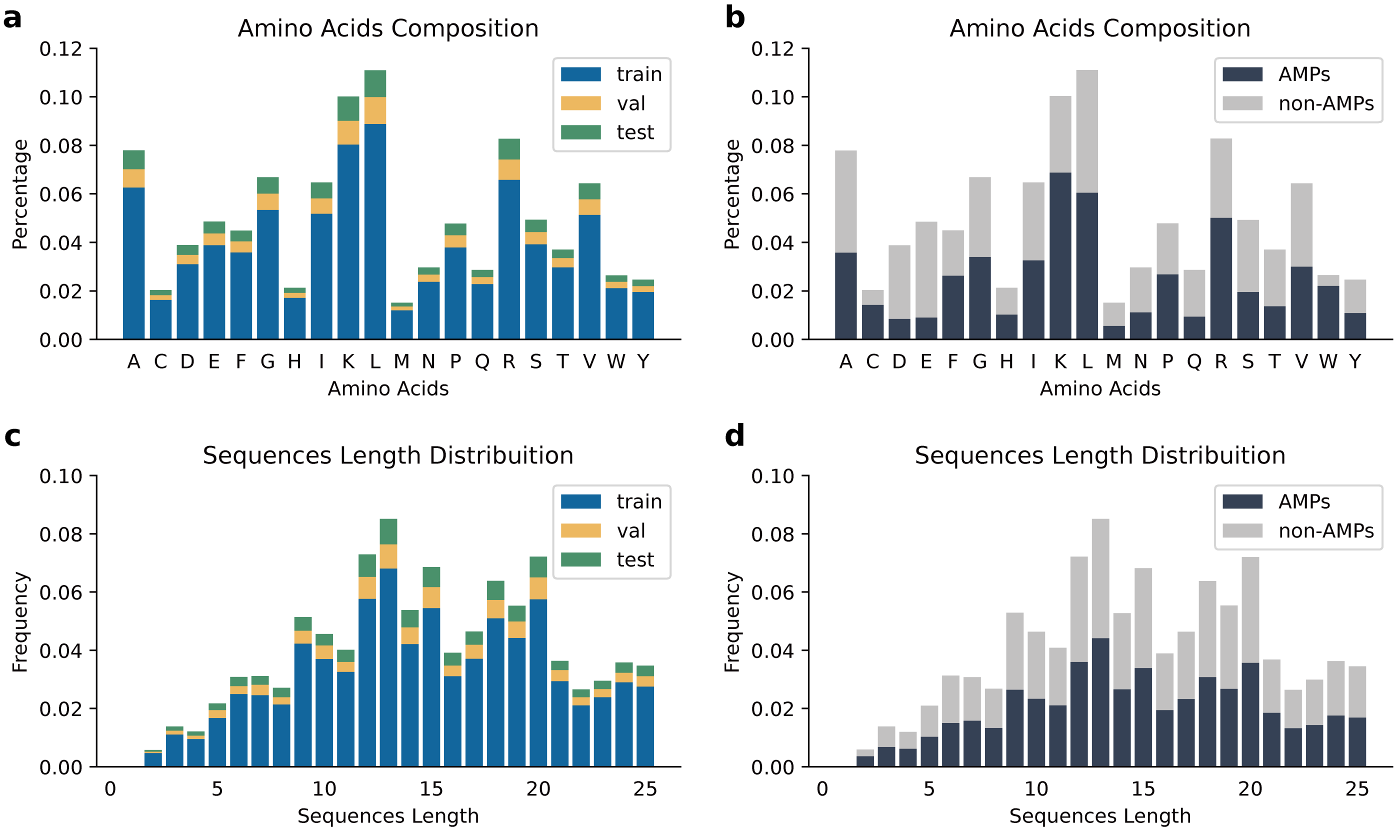


**Supplementary Fig. 1** Amino acid composition and length distribution of the AMP and non-AMP dataset. **a-b** Distribution of amino acid composition divided by **a** training/validation/test set and **b** AMP/non-AMP. **c-d** Sequence length distribution divided by **c** training/validation/test set and by **d** AMP/non-AMP.

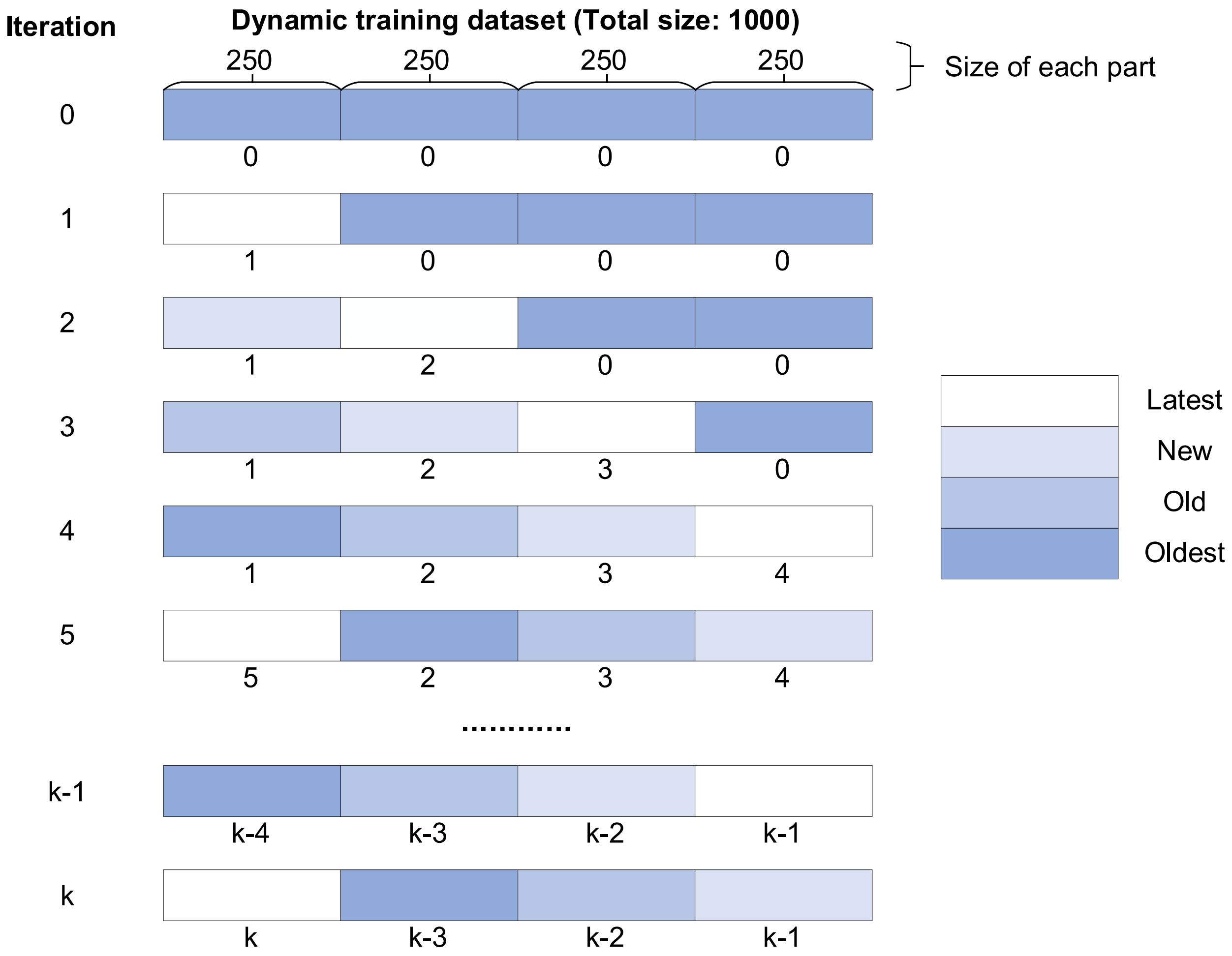


**Supplementary Fig. 2** The real-time knowledge-updating strategy for the dynamic dataset. Each row shows the state of the dataset after each iteration of updates. The number below each set of data represents the iteration in which the set was updated.

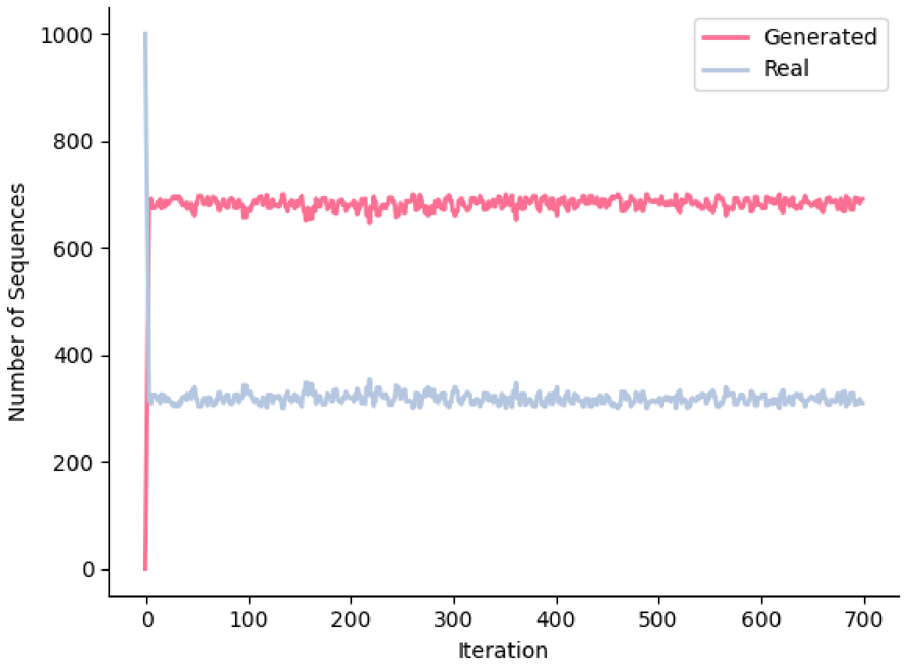


**Supplementary Fig. 3** Changes in the composition of the dynamic training dataset during the MPO stage.

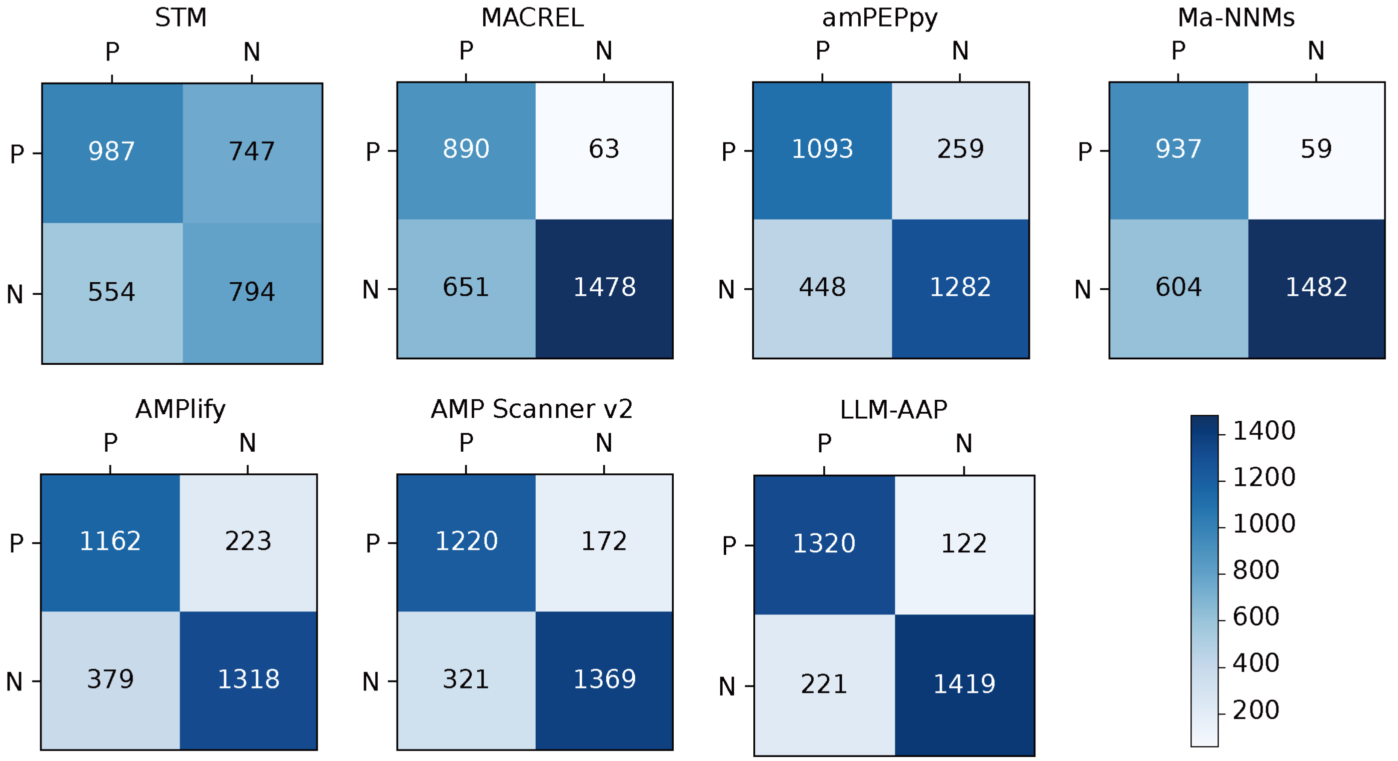


**Supplementary Fig. 4** Confusion matrix of methods for predicting AMPs identification on the independent test set. The x-axis represents the true label, and the y-axis represents the predicted label.

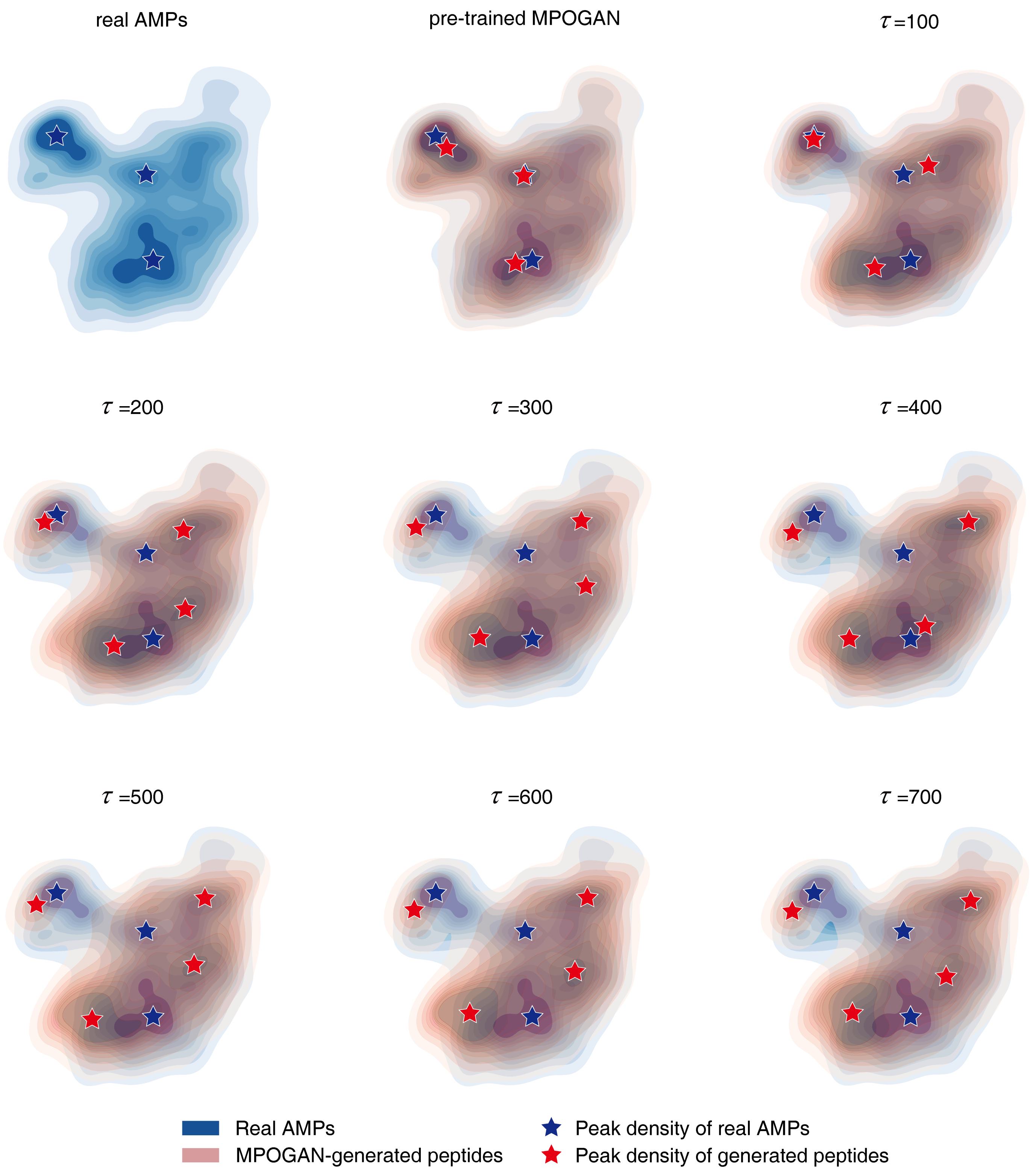


**Supplementary Fig. 5** Visualization of AMPs dataset (real AMPs) and generated peptides from different training stages using Uniform Manifold Approximation and Projection (UMAP) kernel density plot. The data comprises the real AMPs dataset (15406 sequences), peptides generated by the pre-trained MPOGAN (5000 sequences), and peptides generated at various iterations ($\tau$) in the MPO stage (100, 200, 300, 400,

500, 600, and 700 epochs, each with 5000 sequences). Nine physicochemical properties (charge, isoelectric point (pI), aromaticity, Eisenberg hydrophobicity, hydrophobic moment, hydrophobic ratio, charge density, instability index, and aliphatic index) are calculated for all sequences using modlAMP and then standardized. Subsequently, these properties are reduced to a common two-dimensional space. In the visualization, the AMPs dataset is represented in blue, while peptides generated at different stages are in red, with darker regions indicating higher data density. Peaks of data density for the AMPs dataset and generated peptides are denoted by blue and red pentagrams, respectively. It is observed that throughout the training process, the generated peptides exhibit a similar data manifold to the real AMPs. Moreover, the pre-trained MPOGAN effectively captures the density distribution of the AMPs dataset, demonstrating the effectiveness of the pre-training stage. During the MPO stage, the peak of data density for generated sequences gradually shifts, stabilizing after ~500 iterations, indicating the convergence of the MPO process. In conclusion, MPO-GAN is able to generate peptides resembling the true AMPs distribution but exhibiting preference differences.

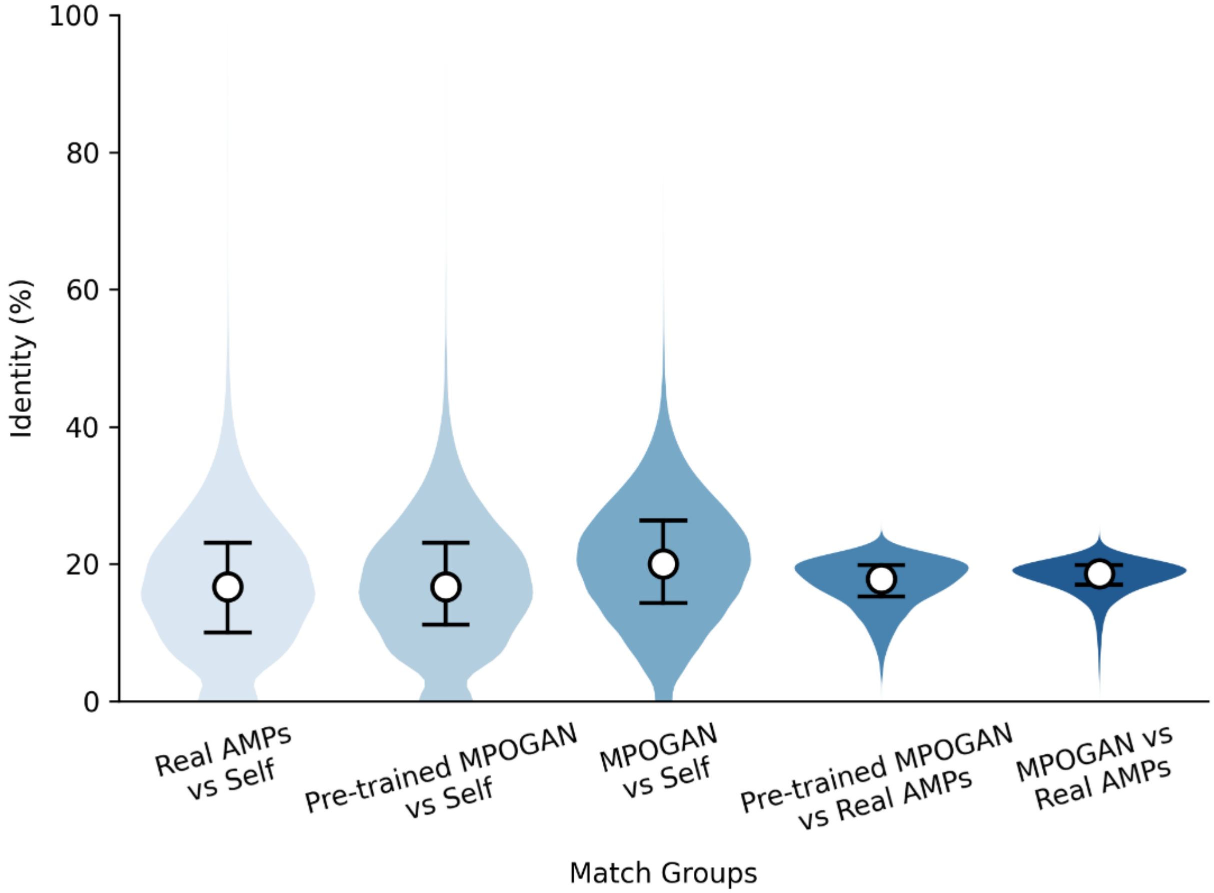


**Supplementary Fig. 6** Violin plots showing distributions of match scores obtained from comparisons between different groups of peptides. Identity is obtained through pairwise sequence alignment of amino acid sequences. The white dots mark the median of each distribution, and the black horizontal lines denote the interquartile range of each distribution.

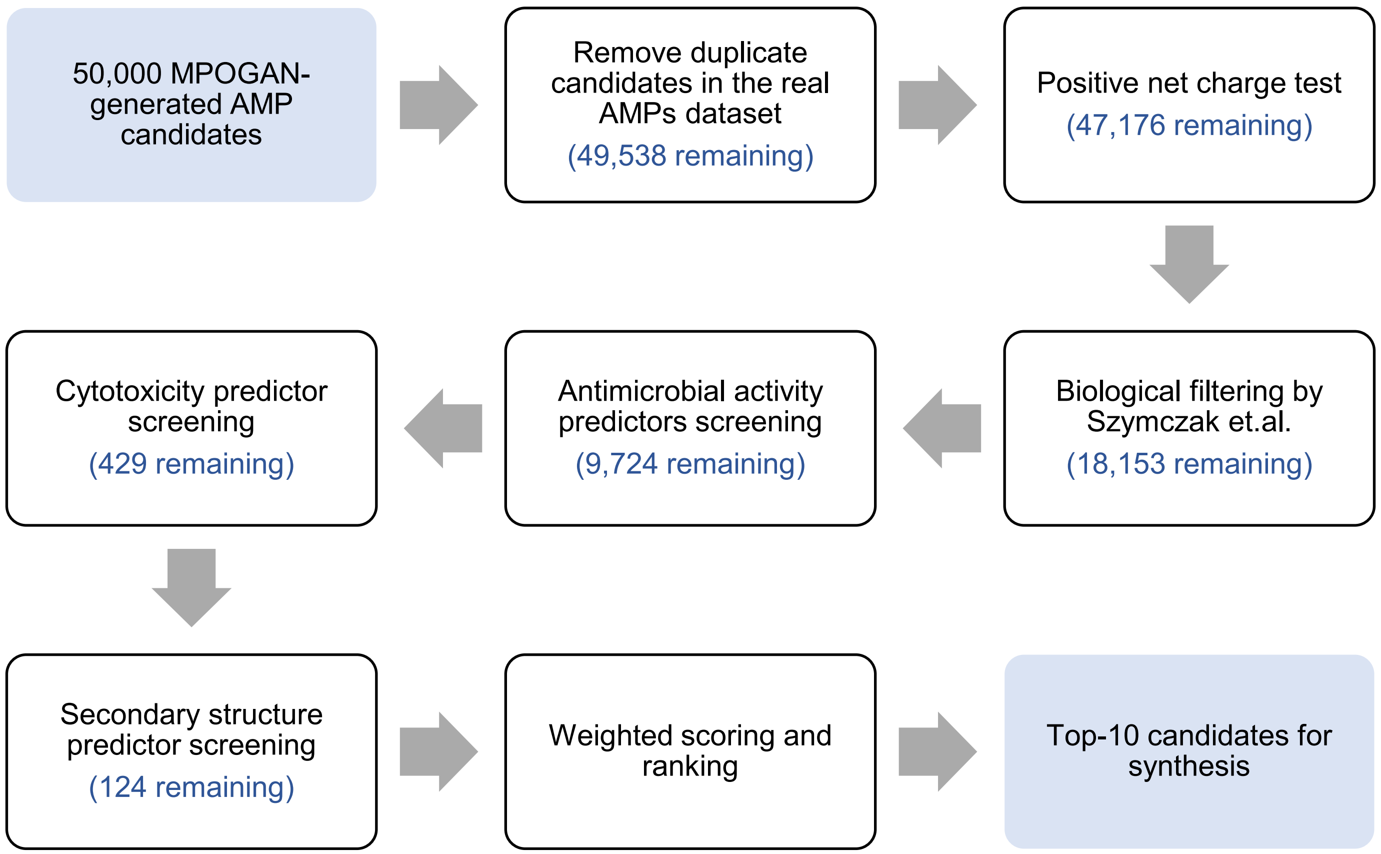


**Supplementary Fig. 7** Preliminary selection process before wet-experiment validation.

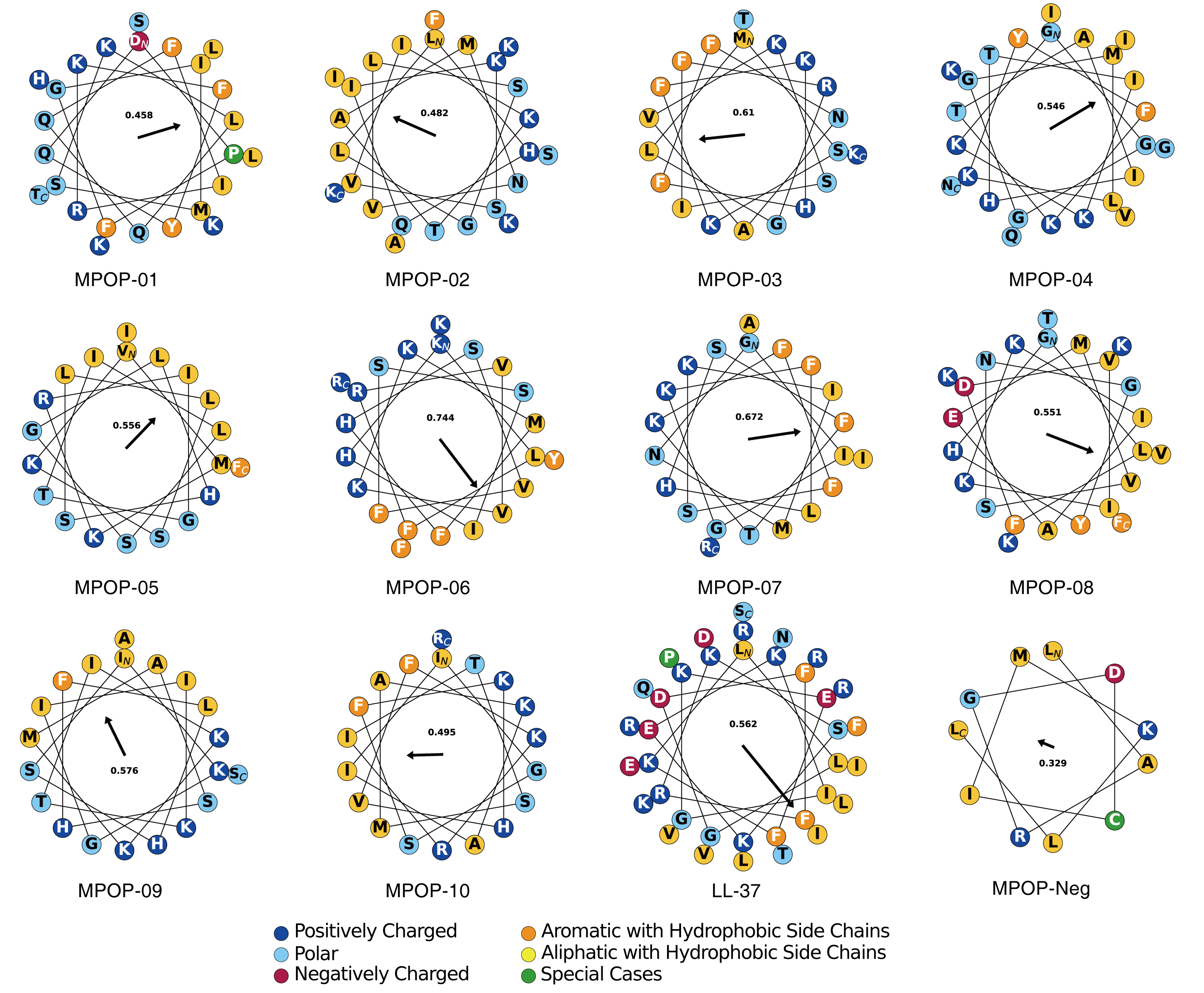


**Supplementary Fig. 8** Alpha-helical wheels of 12 synthesized peptides.

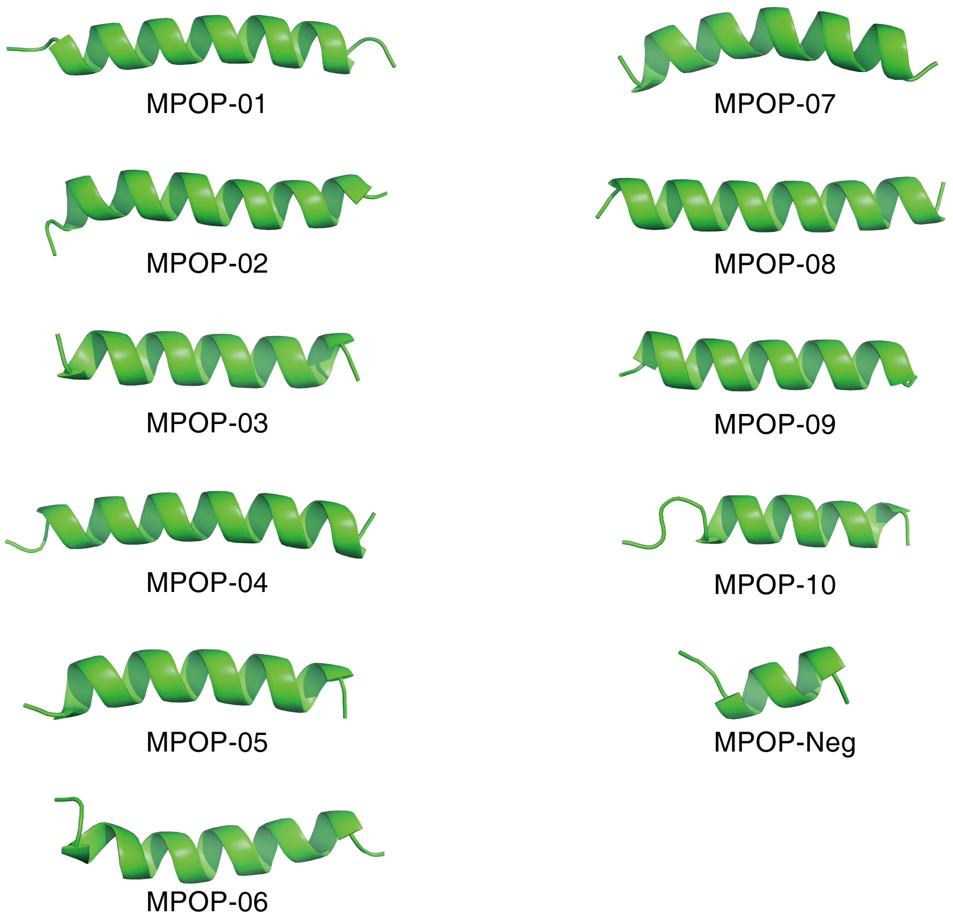


**Supplementary Fig. 9** 3D structures of synthesized peptides predicted using AlphaFold3.

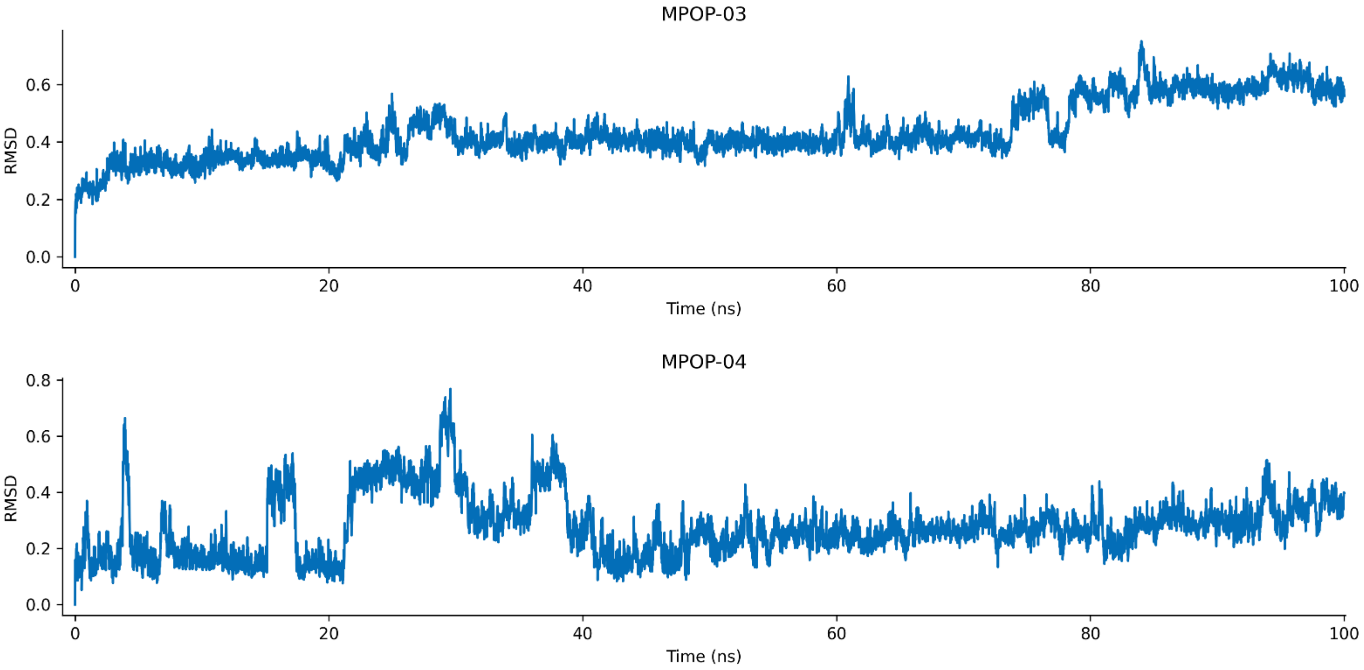


**Supplementary Fig. 10** RMSD for molecular dynamics simulation of MPOP-03 and MPOP-04.
